# CXCR2-mediated recruitment of adaptive NK cells with NKG2C/HLA-E dependent antigen-specific memory enhances tumor killing in ovarian cancer

**DOI:** 10.1101/2024.03.28.585607

**Authors:** Yizhe Sun, Andrea Rodgers-Furones, Okan Gultekin, Shruti Khare, Shi Yong Neo, Wenyang Shi, Lidia Moyano Galceran, Kong-Peng Lam, Ramanuj Dasgupta, Jonas Fuxe, Sahar Salehi, Kaisa Lehti, Dhifaf Sarhan

## Abstract

Natural killer (NK) cells have emerged as promising effectors in cancer immunotherapy due to their ability to recognize and eliminate tumor cells. To investigate the immunological memory and tumor reactivity of adaptive (a)NK cells in the context of desmoplastic tumors, we used human ovarian cancer as a model. Through *in vitro* culture systems resembling dendritic cell (DC)-mediated T cell activation, we demonstrated that aNK cells exhibit antigen-specific cytotoxic responses and memory generation towards ovarian tumor antigens. Furthermore, mature DCs presenting tumor-associated antigens induced the expansion of aNK cells, suggesting antigen-specific proliferation. Single-cell transcriptomics revealed a distinct genetic signature of aNK cells in tumor samples, characterized by a cytotoxic phenotype and interactions with myeloid cells, particularly DCs. The spatial analysis confirmed the intratumoral presence of aNK cells, with higher abundance in the tumor nest compared to conventional (c)NK cells. Functional assays demonstrated the cytotoxicity of expanded aNK cells against autologous ovarian tumors, accompanied by an activated receptor profile. Importantly, aNK cells displayed antigen-specific memory responses towards primary tumors, maintaining specificity over time. Blockade of NKG2C and HLA-E influenced aNK cell recall responses, indicating their roles in the adaptive NK cell immune memory. Additionally, CXCR2 was essential for efficient aNK cell migration toward tumors. These findings shed light on the therapeutic potential of aNK cells in ovarian cancer immunotherapy, highlighting their ability to develop immunological memory and effectively eradicate tumor cells.

## Introduction

Despite current cancer immunotherapies having improved survival rates, clinical outcomes, and treatment responses, only 20-40% of patients benefit and present prolonged remission. Numerous studies have delineated the intricate interplay between tumor cells and the tumor microenvironment (TME), and their interactions affect treatment outcomes, limiting the immunotherapy efficacy. Key factors encompass the deficiency in antigen presentation and the restricted infiltration of cytotoxic immune effector cells, such as CD8 T cells and Natural Killer (NK) cells, into the tumor (*1–6*). NK cells have traditionally been viewed as the primary effectors of innate immunity, lacking antigen specificity and memory (*7*). However, recent evidence indicates the existence of a subset of NK cells, so-called adaptive (a)NK cells, with adaptive immune features. These include memory-like properties, long-term persistence, substantial preferential expansion in response to viral infection, and heightened antibody-dependent effector functions (*8–11*). We have previously demonstrated that aNK cells can establish memory responses to viral peptides following dendritic cell (DC) priming. Certainly, these aNK cells exhibited the ability to initiate recall responses upon secondary stimulation with viable, virally infected cells, distinguishing them from conventional (c)NK cells (*12*). In this study, we sought to investigate whether aNK cells can establish antigen-specific memory responses to tumors and elucidate their role in desmoplastic tumors as a potential immunotherapy agent. High grade serous ovarian cancer (HGSOC) is ideally suited to the study of aNK cells especially where NK cells have been shown to express the NK cell checkpoint receptor NKG2A (*13*). HGSOC has high TME heterogeneity owing to its clinical presentation with multisite abdominal disease and standardized treatment often with surgical debulking enabling *in situ* and *ex vivo* investigations. Moreover, HGSOC classification as a moderate immune-hot tumor increases the probability of encountering tumor infiltrating aNK cells. aNK cells with their inherent ability to resist the TME-mediated immune-evading mechanisms (*14, 15*) and the capacity to acquire immune memory and mount recall responses, render them attractive targets for anti-tumoral immunotherapeutic strategies. For this reason, it is of significance interest to investigate their role in cancer and gain novel insights into the mechanisms that govern their adaptive-like cytotoxic properties. Of note, aNK cell role in desmoplastic tumors is barely known and there is a limited knowledge about their immunological memory to tumors.

Here, our *in vitro* and *ex vivo* investigations indeed revealed a remarkable capacity of aNK cells to recognize autologous tumors, establish memory, and infiltrate the tumor nest by mechanisms involving HLA-E, NKG2C, and CXCR2. Notably, aNK cells obtained from human ovarian biopsies demonstrated superior recognition and killing of autologous tumor cells compared to cNK cells. Moreover, our data underscore the pivotal role of DC in enhancing aNK cell performance. Collectively, our findings represent, to our knowledge, give first insights into the adaptive-like properties of aNK cells to form memory responses to desmoplastic tumors and such specific cytotoxicity is crucial for ovarian cancer cell killing.

## Materials and methods

### Antibodies and reagents

All antibodies and chemicals included in this study have been listed in the **Key Resources Table**.

### Study approval

The HGSOC tumor samples were collected from the patients participating in a Phase III clinical trial (Intra-Peritoneal Local Anesthetics in Ovarian Cancer trial (IPLA-OVCA) (https://clinicaltrials.gov/ct2/show/NCT04065009). Written and informed consents were obtained from all patients before inclusion in the trial in accordance with the Declaration of Helsinki. The protocol for patient participation was approved by the local Ethics Committee and the Institutional Review Board (Dn°. 2019-05149 and 2015/1862-32).

### Cell lines and primary tumor cells

TYK-nu (Japanese Collection of Research Bioresources Cell Bank, Osaka, Japan) and CFPAC (a pancreatic cell line, the American Type Culture Collection), were cultured in MEMα, or RPMI medium respectively (Thermo Fisher Scientific) supplemented with 10% heat-inactivated fetal bovine serum (FBS) (Thermo Fisher), 1% penicillin/streptomycin (Nordic Biolabs), referred to complete medium (CM). The CFPAC cell line was maintained in RPMI CM. Lastly, the K562 cells transfected with HLA-E*0101 (kindly obtained from Dr. Michaëlesson’s laboratory, Karolinska Institutet) were grown in RPMI CM supplemented with 200 µg/mL geneticin (Thermo Fisher) for continued HLA-E*0101 cell selection. All cell lines were tested negative for mycoplasma contamination using the Mycoalert™ Mycoplasma Detection Kit (Lonza Bioscience Solutions) prior to use in functional assays.

Human primary ovarian tumor cells were derived from IPLA patient tumor biopsies. Upon arrival, the tumor specimens were divided into two fragments: one for tumor-infiltrating NK cell expansion and the other for primary tumor cell isolation. The latter underwent full digestion in harvest medium (RPMI medium with 100ug/mL RNase I and 150ug/mL Liberase) for 30 minutes at 37 ℃. After digestion and red blood cell lysis, the digested tumor tissues were finely filtered through a 70μm cell strainer. The tumor cell suspension was then transferred to a T75 tissue culture flask with RPMI medium and 10% patient-derived ascites, with cell outgrowth monitored weekly. To obtain a primary cell culture devoid of tumor-associated cells like myeloid cells, adipocytes, fibroblasts, or endothelial cells, the tumor cells were harvested and isolated using the Tumor Cell Isolation Kit once reaching 90% confluency, following the manufacturer’s instructions. Tumor cell purity was assessed through the expression of the tumor/epithelial marker pan Cytokeratin (PanCK) and the absence of the fibroblast marker S100A4 expression. To eliminate or inhibit potential mycoplasma contamination, 1 µg/mL ciprofloxacin hydrochloride was added throughout the tumor culture. Cells were cryopreserved with 90% FBS and 10% Dimethyl Sulfoxide (DMSO) at each cell passaging.

### Tumor cell lysate preparation

The TYK-nu and IPLA tumor cells were harvested, washed with phosphate-buffered saline (PBS), and resuspended at a concentration of 10^7^ cells/mL in PBS. Subsequently, cells underwent lysis through six consecutive freeze-thaw cycles using liquid nitrogen and a 56°C water bath. The resulting lysate was filtered through a 70 µM cell strainer (Falcon). Finally, protein concentration was determined using a NanoDropTM Protein A280 (Thermo Fischer Scientific), and the lysates were stored at -20°C until use.

### Isolation and sorting of aNK and cNK cells

For the microbead-based isolation of aNK and cNK cells, we initially isolated bulk NK cells following the manufacturer’s guidelines (Miltenyi Biotec). Subsequently, the isolated NK cells were stained with PE-conjugated anti-NKG2C antibody. NKG2C+ NK cells were then selectively isolated as aNK cells using anti-PE microbeads (Miltenyi Biotec), while the flow-through, representing unlabeled NK cells, was collected as cNK cells. The buffer used throughout the process was prepared with sterile PBS at pH 7.2, containing 0.5% FBS and 2mM EDTA (Thermo Fisher Scientific). For the FACS sorting of aNK and cNK cells, bead-isolated NK cells were initially stained with BV510 anti-human CD3, PE-Cy7 anti-human CD56, APC anti-human CD57, and PE anti-human NKG2C antibodies. Sorting was conducted on a BD FACSARIA III cytometer at the Flow Cytometry facility (Huddinge, Karolinska Institutet). Cells characterized as CD3-CD56+CD57+NKG2C+ were identified as aNK cells, while CD3-CD56+CD57+NKG2C-cells were classified as cNK cells.

### Autologous dendritic cell (DC)-NK cell co-cultures

Peripheral blood mononuclear cells (PBMCs) were isolated from healthy blood donors (provided by the Stockholm blood bank) through Ficoll-Hypaque (GE Healthcare) density-gradient centrifugation. The aNK frequency (CD56+CD3-CD57+NKG2C+) in each donor was assessed prior to bead-isolation using a CytoFLEX cytometer (Beckman Coulter Life Sciences), and only donors with ≥6% aNK were included in the study.

Monocytes were isolated from PBMCs using CD14 microbeads (positive selection), respectively, following the manufacturer’s instructions. Monocytes were seeded at 1 x 10^6^ cells/mL in CellGro serum-free DC medium (CellGenix) supplemented with 2% human AB serum and 1% penicillin/streptomycin. Differentiation into monocyte-derived dendritic cells (DC) was induced by adding GM-CSF (100 ng/mL, Peprotech) and IL-4 (20 ng/mL, Peprotech) for 48 hours. Immature DCs were then washed, and maturation was induced with specific factors. Immature DCs were also pulsed with tumor antigens by adding 30 µg tumor cell lysate (TYK-nu or IPLA) per one million DCs. DC-NK cell co-cultures were performed in round-bottom 96-well plates with RPMI CM supplemented with 10 ng/mL rhIL-15. The cultures were maintained for one to four weeks, with media replenished every 7 days for co-cultures lasting 2-4 weeks. At the 2nd, 3rd, and 4th weeks, NK cells were restimulated with autologous IPLA and TYK-nu cells or allogenic tumor cells (CFPAC) at an E:T ratio of 1:1. Following restimulation, immune staining was performed. In additional experiments, blocking antibodies (5 μg/ml) to anti-NKG2C (clone 134591), anti-NKG2A (clone 131411), or anti-HLA-E (clone 3D12), anti-MHC I (clone W6/32), and MHC II (clone M5/114.15.2), or a control isotype-matched antibody IgG (clone Poly4053) (BioLegend, R&D systems) were added at the primary phase (day 0 and 7) or at the secondary restimulation phase (at day 21, during 6h stimulation).

### Ex vivo expansion of tumor-infiltrating NK cells

Tumor specimens, dissected into 1-3 mm^3^ fragments, were cultured in 24-well plates, excluding adipose tissue and necrotic areas. Tumor-infiltrating lymphocytes (TILs) medium consisting of RPMI supplemented with 6,000 IU/mL IL-2, 5 ng/mL IL-15, and 10% human AB serum supported lymphocytic outgrowth (TILs CM). Medium replacement occurred every 24-48 hours, and visible lymphocyte growth prompted a CM change within a week. Seven to ten days into culture, 80% of the medium was substituted with CM plus 10% human AB serum, 2,000 IU/mL IL-2, and 5 ng/mL IL-15. Confluent wells were split into daughter wells to maintain a cell density of 0.8-1.6 x 10^6^ cells/mL. Each well represented a distinct TIL clone. After 12-14 days, cNK and aNK cell frequencies from each TIL clone were quantified using a CytoFLEX cytometer. Clones with ≥10% CD45+ NK cells or ≥5% NK cells were harvested. TIL clones with similar CD45+, NK cells, and T cells patterns were pooled. NK cell-derived TIL clones underwent a rapid expansion protocol. Cells were cultured in RPMI1640 medium with 1% penicillin/streptomycin, 10% human AB-serum, 200 IU/mL rh-IL2, 5 ng/mL rhIL-15, and 100 ng/mL IL-21. Co-culture with allogeneic irradiated feeder cells (K562-HLA-E*0101 at 5:1 ratio) continued for four weeks, maintaining cell densities around 1 × 10^6^ cells/mL. Following assessments, the expanded TILs were cryopreserved with 90% FBS and 10% DMSO.

### TIL phenotyping

For the assessment of one-month-expanded TIL phenotype from ovarian tumor biopsies pre- and post-coculture with tumor cells, TILs were cocultured with tumor cells for 6 hours. Subsequently, cells were stained for NK cell markers as detailed earlier. Following staining, cells were fixed and permeabilized using the eBioscience Foxp3/Transcription factor staining buffer set according to the manufacturer’s instructions. Intracellular staining was then conducted at room temperature in the dark for 40 minutes with anti-FCεR1γ. Samples were acquired with a CytoFLEX cytometer, and FlowJo v.10.5 software (BD Biosciences) was utilized for analysis.

### Flow cytometry-based killing assay

To examine the killing capacity of the TILs against autologous cancer cells, the TILs were co-cultured 6 hours with the target tumor cell (IPLA) or an irrelevant tumor cell TYK-nu at effector: target (E:T) ratio of 1:1. The cells were collected, and stained with anti-CD56, anti-CD45, Live/Dead cell fixable dye and anti-CD3. Cells were then fixed with 4% paraformaldehyde (see above in the TIL phenotyping section), washed, and treated with trypsin to detach tumor cells. Lastly, cells were washed, filtered and analyzed by flow cytometry.

### Analysis of cytokine production and degranulation by flow cytometry

Pro-inflammatory cytokine production and degranulation were assessed in DC-NK cell and TIL cultures using flow cytometry. Following the DC-NK cell co-culture, NK cells were restimulated with viable tumor cells presented by DCs (IPLA or TYK-nu) or an irrelevant tumor cell line (CFPAC) at an E:T ratio of 1:1. Likewise, TIL cultures were co-cultured with the target tumor cell (IPLA) at an E:T ratio of 1:1. For both, restimulation occurred in RPMI1640 medium containing human CD107a antibody, along with protein transport inhibitors Golgiplug (BD Biosciences) (1:1000) and Golgistop (BD Biosciences) (1:1000) for 6 hours prior to staining. Following surface staining, cells were fixed and permeabilized using the eBioscience FOXP3/Transcription Factor Fixation/Permeabilization buffer (Thermo Fisher Scientific, USA) for 20 minutes at room temperature in the dark. Subsequently, cells were stained for intracellular markers, including FCεR1γ, anti-Ki67, anti-IFNγ, and anti-TNFα. After washing, cells were acquired with a CytoFLEX cytometer and analyzed using FlowJo.

### Migration assay

Following three weeks of NK-DC coculture, aNK and cNK cell migration was evaluated in transwell assays (5 µm) allowing for only active migration of NK cells, laid onto the upper insert, toward the bottom where indicated target was cultured. In a few experiments, the migration of TIL aNK and cNK cell migration was evaluated toward autologous tumors in the presence of control IgG antibody or blocking antibody to CXCR2 (5 µg/mL) during six-hour assay.

### Multiplexed Tissue Staining and Image Acquisition

Paraffin-embedded ovarian tumor tissue slides underwent a two-phase baking process: first, at 37°C overnight, followed by a second phase at 60°C for one hour. Subsequent staining was performed using the BOND RXm Automated IHC Stainer (Leica Biosystem), adhering to the following staining sequence: FcεRγ (1:200, Opal 520), CD3 and CD11c (1:500 and 1:100 respectively, Opal 540), CD56 (1:50, Opal 570), CD57 (1:100, Opal 620), NKG2C (1:100, Opal 650), and a combination of two pan-Cytokeratin antibodies, AE1/AE3 and C11, (1:50 and 1:250 respectively, Opal 690). Antigen retrieval for FcεRγ, CD3/CD11c, CD56, NKG2C, and Pan-CK was performed using BOND Epitope Retrieval Solution 1 (citrate-based), while CD57 antigen retrieval needed BOND Epitope Retrieval Solution 2 (EDTA-based), both facilitating heat-induced epitope retrieval. Following staining, slides were preserved with ProLong™ Diamond Antifade Mountant (Thermo Fisher Scientific). The Vectra 3 Automated Quantitative Pathology Imaging System was employed to capture images of the tissue sections. Comprehensive analysis, including unmixing, tissue segmentation, cell segmentation, and cell phenotyping, was conducted using inForm 2.6.0 software. This allowed for the extraction of cell counts, distribution, and neighborhood analysis by using phenoptrReports (Akoya Biosciences). Finally, the phenotyping maps were generated using CytoMAP (*16*), extracting the data obtained from inForm.

### scRNAseq data analysis

Raw gene counts were downloaded from GEO (GSE184880 (*17*)) and analyzed using the Seurat package (version 4.4.0) (*18*). Seurat includes functions to calculate quality control metrics, dimensionality reduction, clustering, differential gene expression analysis and data visualisation. Customized filtering criteria were used for each sample to retain sufficient cells for further analysis. Cells with fewer than 200 genes and more than 20 or 25% mitochondrial RNA were excluded. Cells with more than 7500, 9000 or 10000 genes were excluded to remove possible doublets. Additional doublet filtering was performed using scDblFinder (*19*). After filtering, the data was subjected to log normalization, followed by scaling and dimensionality reduction using PCA. Graph based clustering was performed using the computed principal components (PCs) and visualized using UMAP. Varying number of PCs were employed for clustering/subclustering to obtain well defined clusters. Cells were annotated based on gene expression signatures computed using differential gene expression analysis and literature-based evidence.

### Interaction analysis based on scRNAseq data

CellphoneDB (*20*) was employed to compute interactions between myeloid and NK clusters by utilizing its database of known ligand-receptor interactions. This tool assessed the co-expression of ligand-receptor pairs in the single-cell transcriptomic data. The results were refined by filtering putative interactions with a p-value < 0.01 and requiring receptor and ligand expressions to be > 0.5, thus retaining the most significant interactions. Subsequently, a heatmap illustrating the number of interactions between myeloid clusters and various NK clusters was generated using ggplot2 in R (version 4.3.1). Furthermore, a chord diagram depicting interactions was created, focusing on cDC1 and cDC2 as the source and NK clusters as targets. In this case, a receptor and ligand expression cutoff of 0.1 was applied, and interactions involving protein multimers were excluded for clarity and accuracy.

### Statistics Analysis

All experiments were repeated independently at least three times and one representative and accumulative data are presented. All numeric data were subjected to normal distribution using the Shapiro-Wilk test and QQ plots tests before further statistical analysis. For the comparison within groups, we used the multiple paired T test without considering the effects within groups or parametric two-way ANOVA. Student T-test was used when comparing two groups. Single linear regression and correlation analyses were performed. All statistical tests were two-sided. All p-values from multiple comparisons were corrected by using the FDR method <0.05 and all non-significant p-values are not indecated. The Prism v9.2 software (GraphPad) was used for statistical analyses.

### Data availability statement

The codes and datasets are available in the GitHub repository: https://github.com/shrutikhare-git/scRNAseq_analysis_adaptive_NKs

## Results

### aNK cells exhibit immunological memory towards ovarian tumor antigens

Recently, we demonstrated that aNK cells, when primed by viral peptide-loaded DCs, exhibit antigen-specific responses with the capability of immune memory generation (*12*). In this study, we sought to examine the hypothesis that aNK cells develop antigen-specific memory in response to desmoplastic tumors. To test this, an *in vitro* culture system resembling DC-mediated T cell activation was set up (**Supplementary Figure 1A**). Tumor cell lysate was generated from an ovarian cancer cell line TYK-nu, using an adapted clinical protocol for DC-vaccination in melanoma patients (*21*). The responses of aNK and cNK cells (**Supplementary Figure 1B**, Gating strategy) towards the TYK-nu cells were used as a specificity measure, meanwhile, response to an unspecific pancreatic tumor cell line CFPAC was used as a control. Following three weeks of NK-DC coculture, aNK cells displayed an antigen specific response in a secondary simulation towards TYK-nu cells in form of degranulation (CD107a) and cytokine production (IFNγ and TNFα) compared to first time stimulation and insignificant increased response in cNK cells. On the contrary, cNK cells exhibited unspecific inherent anti-tumor activity towards TYK-nu and CFPAC (**Figure 1A, 1B, Supplementary Figure 1C**). These results suggest that aNK cells gain antigen specificity and process memory formation that is detected several weeks following priming. This cellular behavior resembles the initiation of DC-mediated T cell adaptive responses, which are launched after a few weeks of exposure to a specific antigen. While T cells are known to proliferate upon priming by antigen-loaded DCs, we sought to investigate whether aNK cells do the same and expand under such conditions. Mature DCs presenting ovarian tumor-associated antigens from tumor lysate including TYK-nu cell line (hereby referred to DC^lysate^), were cultured together with autologous bulk NK cells for four weeks. aNK cells co-cultured with DCs for four weeks increased their frequency compared to aNK cells alone **(Figure 1C, 1D)**. aNK cells have been hypothesized to be derived from CD56^dim^-mature/terminally differentiating cNK cells (*22*), thus we performed tumor-antigen priming in FACS-sorted aNK or cNK cells. After four weeks of culture, aNK cells barely proliferated from the sorted cNK cells, compared to the sorted aNK cells, which showed a substantial increase especially when primed with DC^lysate^ compared to DC^-^ **(Figure 1E, 1F)**. In addition, we conducted similar proliferation/differentiation assay but this time in bulk NK cells. Still, aNK cells were the only NK subpopulation that persist in long term culture and had antigen-specific expansion (**Supplementary Figure 2**). These results indicate an aNK cell antigen specific expansion mediated by DC^lysate^ rather than differentiation from cNK cells. Because we did not use the intracellular molecule FcεR1γ to discriminate aNK and cNK cells during the FACS-sorting, residual NKG2C expressing cNK cells may remain in sorted aNK population, but they still failed to proliferate.

**Figure 1.**
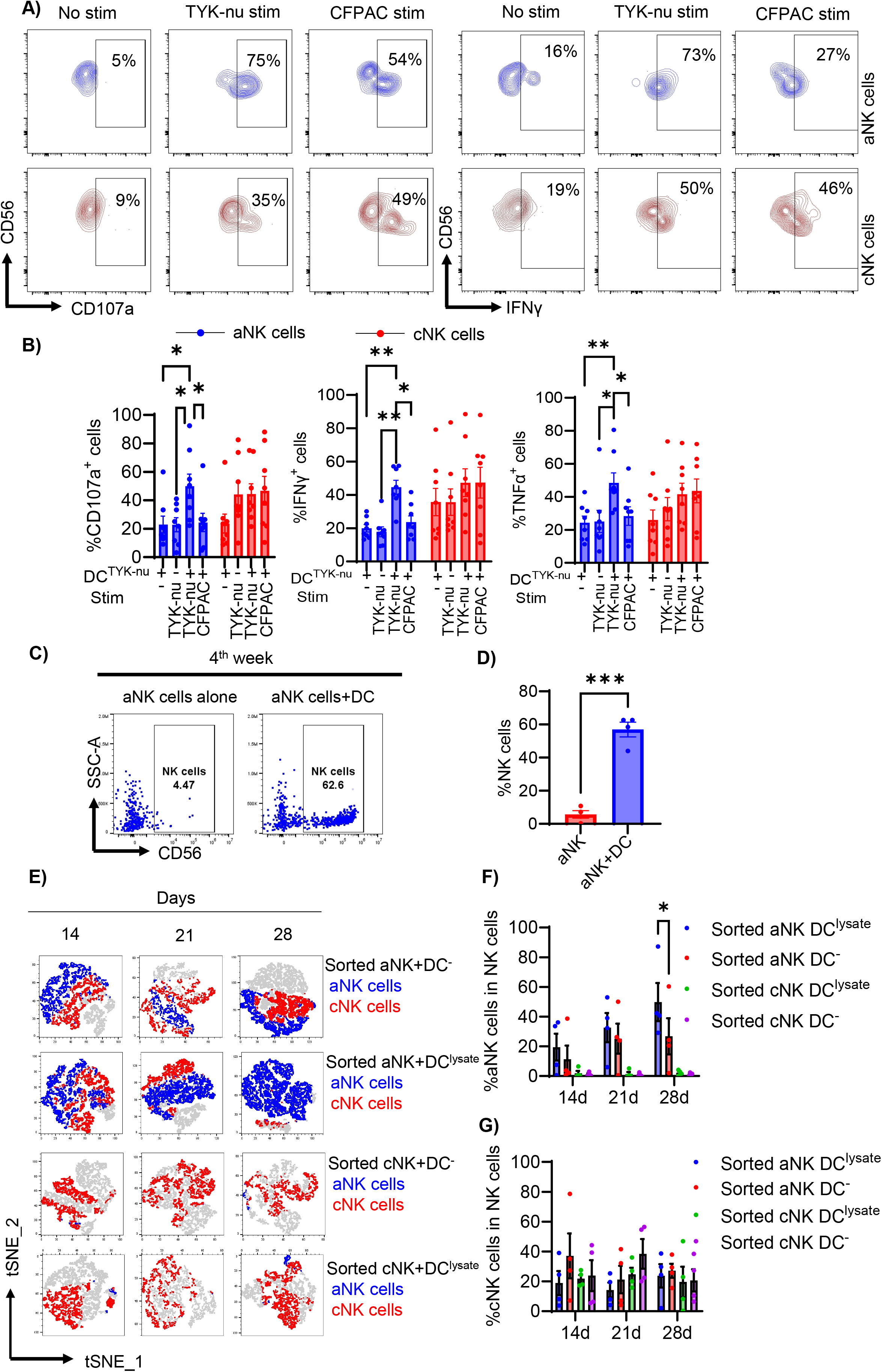
aNK cells present specific immune memory towards established tumor cell line and expansion upon DC coculture with the antigen presentation. A) After three weeks of NK-DC coculture, restimulation with TYK-nu cells was performed. Contour plots show CD107a^+^ and IFNγ^+^ cells among aNK and cNK cells under different conditions: no stimulation after DC coculture, autologous restimulation with TYK-nu, and allogeneic restimulation with CFPAC. **B)** Accumulated data display proportions of CD107a^+^, IFNγ^+^, and TNFα^+^ cells (n=8) across experimental scenarios: priming with DC^TYK-nu^ without restimulation, autologous restimulation with DC^TYK-nu^ + TYK-nu, allogeneic restimulation with DC^TYK-nu^ + CFPAC, and first stimulation with DC^-^ + TYK-nu, for aNK cells (blue) and cNK cells (red). Two-way ANOVA (*p < 0.05, **p < 0.01) was conducted. **C, D)** After four weeks, FACS-sorted aNK and cNK cells were expanded with or without DCs. Paired T test (*p < 0.05, ***p < 0.001) was performed. **E)** tSNE plots illustrate expansions of aNK (blue) and cNK (red) cells over 2 to 4 weeks upon DC coculture with or without tumor lysates. **F-G)** Cumulative frequencies of aNK and cNK cells from two to four weeks upon DC coculture with tumor lysates are shown. Two-way ANOVA (*p < 0.05) was conducted. SEM is shown for cumulative data.

Overall, these data showed that aNK cells recognized tumor antigens previously presented by DCs and acquired immunological memory.

### aNK cell genetic signature in ovarian cancer patients revealed activated cells and interacting with DC

Given the immune suppression resistance of aNK cells in hematological malignancies, we hypothesized that aNK cells will also persist and play role in solid tumors. We first investigated a publicly available ovarian cancer (OC) single cell transcriptomic dataset (GSE184880) (*17*). Raw counts were downloaded from GEO and analyzed using Seurat version 4.4.0 (*18*). By Louvain clustering, we identified 10 subpopulations of NK cells in which we found a distinct cluster 1 with known aNK cell phenotype (NKG2A^-^, NKG2C^+^, CD16^high^, CD57^+^ and FCER1G^low/neg^) and other distinct differentially expressed genes (DEGs) associated with NK cell cytotoxicity including perforin and granzyme B (**Figure 2A, 2B and Supplementary Figure 3A**). Likewise, various myeloid populations with distinct DEGs could be identified including monocytes, DCs and macrophage subsets (**Figure 2C, 2B and Supplementary Figure 3B**). To quantify NK-myeloid cell physical interactions, we applied CellPhoneDB (*20*) that utilized a public repository of known ligand-receptor interactions. Notably, aNK cell like cluster 1 appeared to interact with myeloid cells in general at higher frequencies than the other nine NK cell subclusters (**Figure 2D**). Focusing on DCs, we observed more cDC2-NK cells than cDC1-NK cells interactions. A large portion of NK-DC interactions involved NK cluster 1,2 and 3 which were all CD16^+^, PRF1^+^, GZMB^+^ and NKG2A^-^. However, the interaction of cDC1 was more restricted to aNK cell (**Figure 2E**). Exploring the top ligand-receptor underlying cDC1-aNK cell interaction, identified HLA-E-KLRC2 which may play a role in antigen presentation as was demonstrated in our previous study (*12*). Other significant interactions were revealed including CXCL1-CXCR2, HLA-C-KIR2DL3, CLEC2B-KLRF1, which are involved in chemotaxis, NK cell education, and antigen presentation and cytotoxicity, respectively. Collectively, our single cell analysis here suggested that DC-aNK cell interactions may be influenced by NK cell maturation and the extent of immune suppression. At the same time, these DC-NK interactions are suggestive of a potential cross presentation of antigen to confer specific immune memory activated aNK cells.

**Figure 2.**
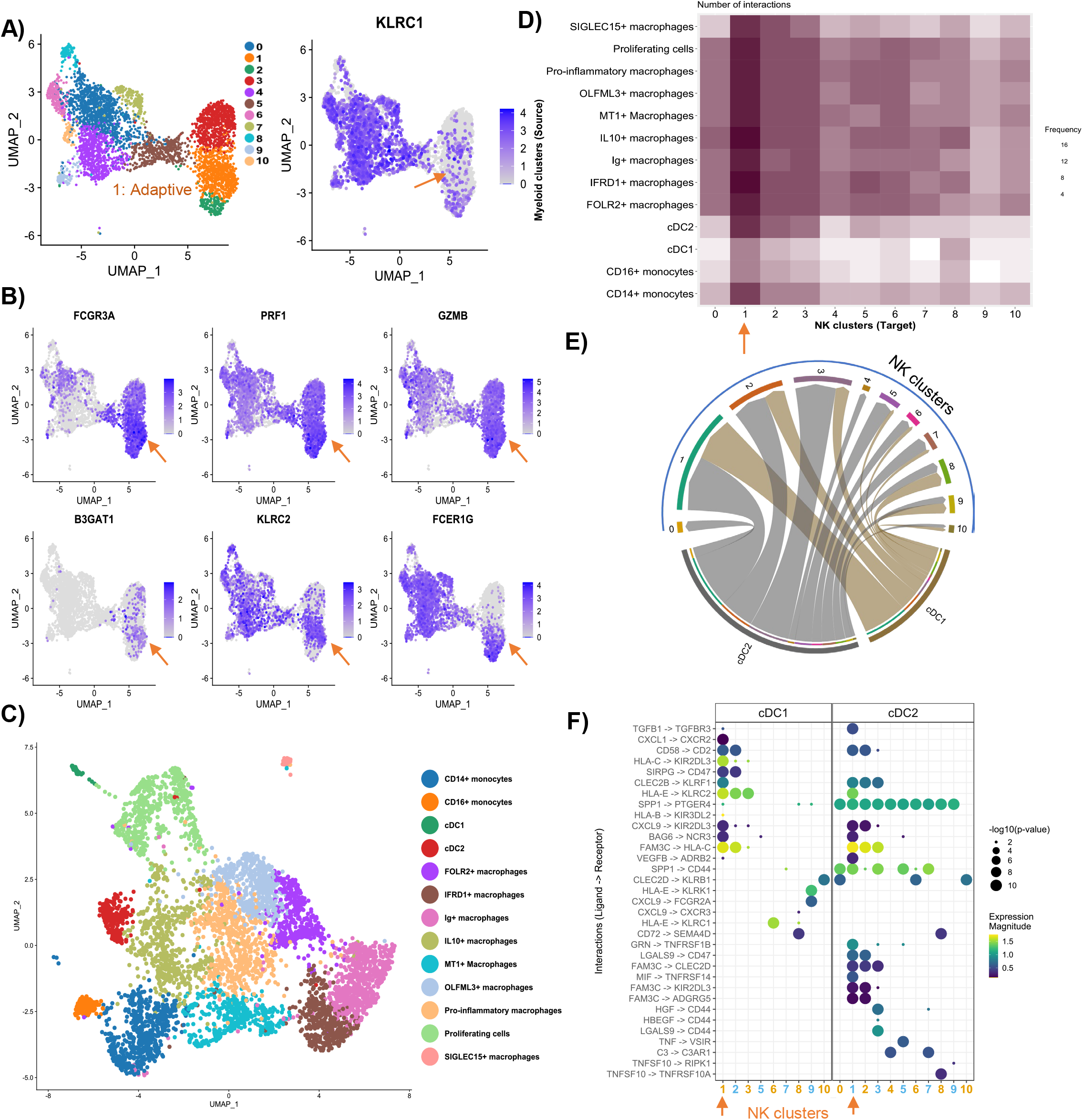
CellPhoneDB analysis reveals a high tendency of adaptive NK cells to engage in DC-NK cell interactions. **(A)** UMAP visualizations display subclusters of NK cells, highlighting specific NK cell markers and **B)** effector genes, with an arrow pointing to the subcluster exhibiting aNK cell characteristics. **C)** A UMAP plot delineates diverse annotated myeloid cell subsets. **D)** A heatmap illustrates the interaction frequencies among various myeloid and NK cell subsets. **E)** A chord diagram depicts the interaction frequencies between different NK cell subsets and DC subsets. **F)** The most significant ligand-receptor pairs facilitating NK cell-DC interactions are listed, with arrows showing connections between the adaptive NK cluster (cluster 1) and two DC subsets, cDC1 and cDC2.

### aNK cells are found in the ovarian intratumoral environment in contrast to cNK cells that were found in the stroma

Next, our aim was to validate the presence of aNK cells in OC and explore their interaction with APCs. Our hypothesis centered on aNK cell infiltration into the TME and their presumed proximity to the tumor nest. Thus, we investigated the presence and distribution of various immune cell populations, including NK cells (CD3-CD56+), T cells (CD3+CD56-), aNK cells (CD3-CD56+CD57+NKG2C+FcεR1γ-), conventional NK cells (cNK cells) (CD3-CD56+CD57+FcεR1γ+), and DCs/antigen presenting cells (APCs) (CD11C), in human HGSOC tissues using Multiplex Immunofluorescence (mIF). Our findings revealed infiltration of NK cells and T cells into ovarian tumors, suggesting an immune-inflamed TME across most specimens studied. Notably, the abundance of tumor-infiltrating aNK cells surpassed that of cNK cells and T cells, indicating their prevalence within the ovarian TME and relative resistance to immune exclusion compared to other NK cell subsets and T cells, who were present in the stroma (**Figure 3A, 3B, 3C**). Further analysis of the spatial relationship between aNK cells and tumor cells revealed closer proximity of aNK cells to tumor cells compared to cNK cells and T cells (**Figure 3D**). Additionally, we examined the distribution of APCs and their proximity to NK cells within HGSOC tissues. Our observations identified APCs presence both within the tumor nest and stroma (**Figure 3E**). Interestingly, aNK cells were found to be closer to DCs in the stromal region, while cNK cells exhibited closer association with APCs within the tumor region (**Figure 3F**). In summary, our results indicate intratumoral presence of aNK cells, which may interact with DCs within the stroma and migrate towards the tumor nest.

**Figure 3.**
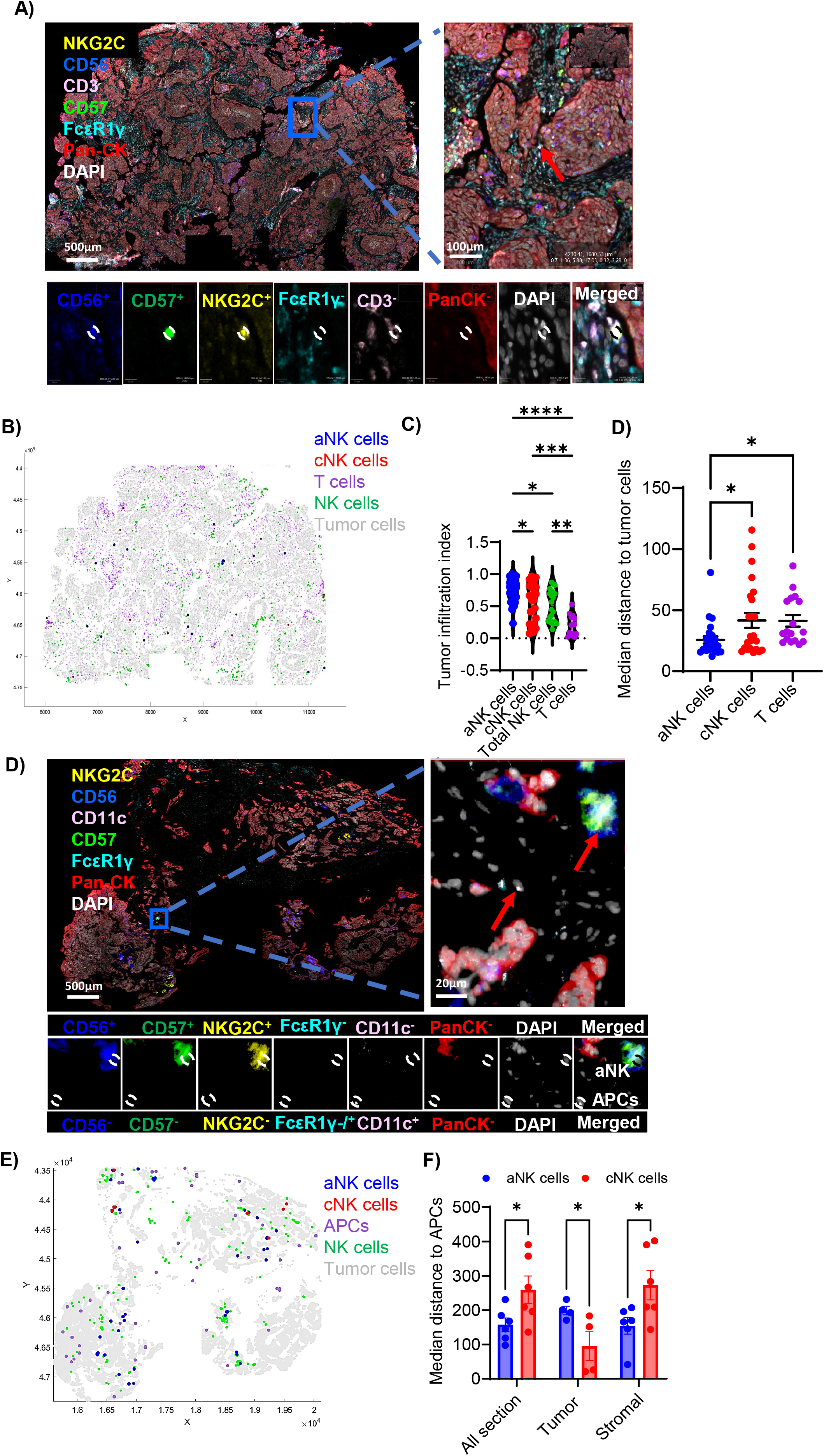
aNK cells are found in closer proximity to ovarian tumor cells and have a more intimate interaction with antigen-presenting cells (APCs) within the stromal region compared to cNK cells. **A)** Representative image demonstrates the identification of aNK cells (CD56^+^CD57^+^NKG2C^+^FcεRγ^-^CD3^-^PanCK^-^). **B)** A phenotyping map shows the distribution of cell types in tumor sections: aNK cells, cNK cells, T cells, NK cells, and tumor cells. **C)** Infiltration rates (left) and median distances (right) of various cell types within tumors. The first analysis showcases the infiltration rates of aNK cells (n=22), cNK cells (n=22), total NK cells (n=22) and T cells (n=14) within tumors through a violin plot, assessing tumor infiltration index. The second analysis evaluates the median distances from aNK cells (n=24), cNK cells (n=24), and T cells (n=17) to tumor cells in the entire sections. A one-way ANOVA test was applied for within-group comparisons. **D)** Representative Image displays the identification of aNK cells and APCs. **E)** A phenotyping map illustrates the distribution of cell types in tumor sections. **F)** Median distances to APCs from NK cells are shown in different tumor areas. It shows the median distance to APCs from aNK cells and cNK cells in tumor area (n=4), stromal area (n=6) and entire section (n=6). Multiple T test was used to the compare within the groups. significance levels: *p < 0.05, **p < 0.01, ***p < 0.001. In this figure, accumulative data are shown as mean ± SEM.

### Tumor-infiltrating aNK cell activity display high expression of activating receptors while cNK cells predominantly express the inhibitory receptor NKG2A

To assess the anti-tumor activity of intratumoral aNK cells within the ovarian TME, we expanded primary NK cells directly from human metastatic ovarian tumor sites for subsequent *ex vivo* functional analysis. Initially, TILs were isolated from millimetrically-dissected tumor specimens and cultured in media supplemented with interleukins IL-2 and IL-15 (**Figure 4A**), known to support NK cell proliferation and survival (*23*). After a two-week culture period, TIL clones that proliferated with ≥10% CD45+ cells and ≥5% NK cells were identified by flow cytometry, combined, and subjected to an expansion protocol. This expansion involved a combination of cytokines IL-2, IL-15, and IL-21 and irradiated feeder K562 cells. Utilization of K562 feeder cells has been demonstrated to mitigate cytokine expansion-induced NK cell senescence, resulting in enhanced yields (*24, 25*). Simultaneously, primary tumor cells were grown to low passage (up to 6) lines. NK cell frequency increased approximately three fold after three weeks of expansion (**Figure 4B**, **4C**). We also found that the frequency of aNK-, cNK-, and T cell was heterogenous among the different TIL clones (**Figure 4E, Supplementary Figure 4A**).

**Figure 4.**
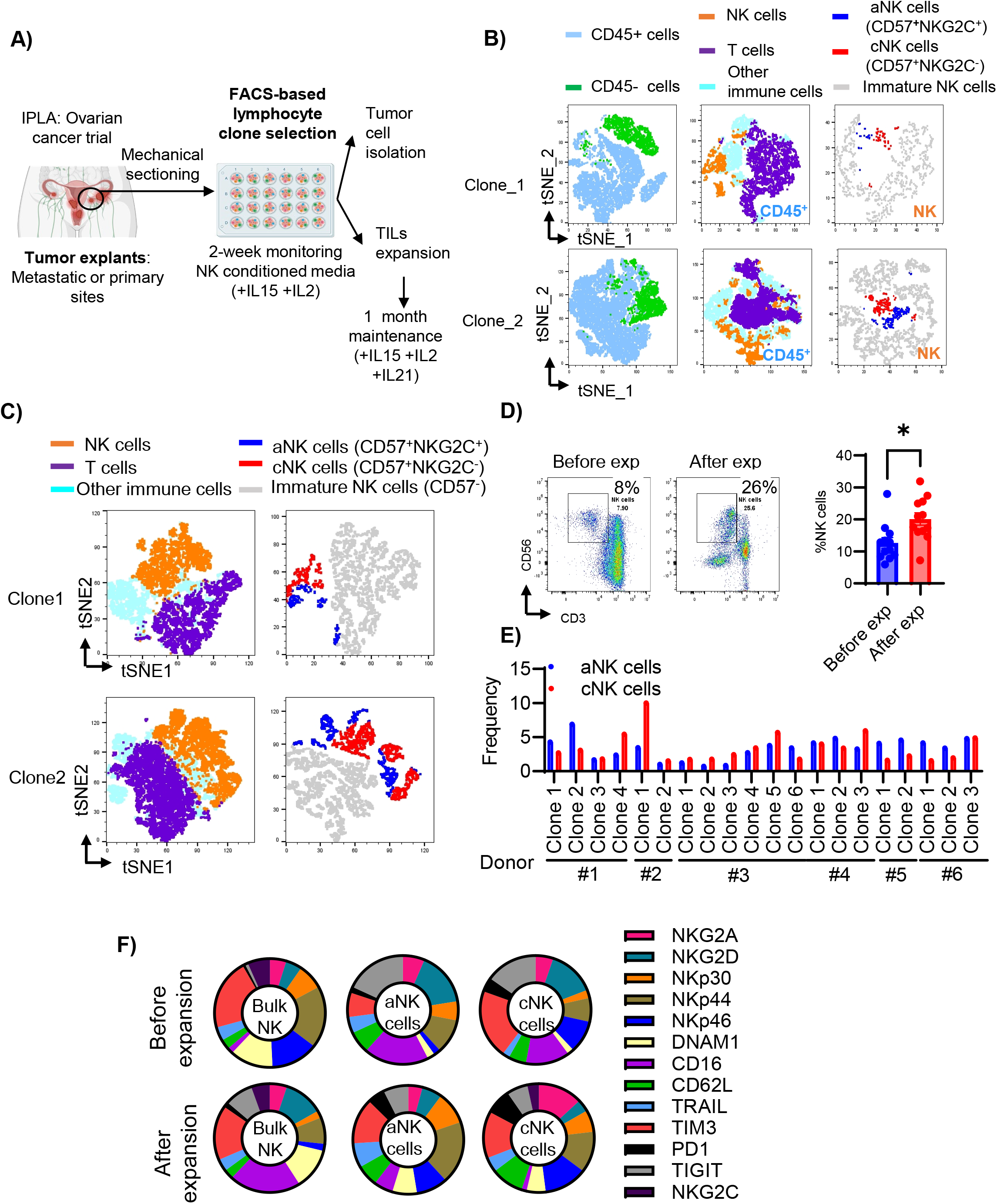
Tumor infiltration lymphocytes (TILs) from ovarian tumors are expanded including aNK cells and cNK cells. **A)** The schematic outline illustrates steps in processing IPLA ovarian tumor samples. **B)** t-SNE visualization shows TIL clone diversity pre-expansion, emphasizing CD45^+^ cells, T cells, and NK cell subsets. **C)** Post-expansion, t-SNE maps illustrate changes in CD45, T cells, and NK cell frequencies. **D)** Flow cytometry analysis comparing NK cell frequencies pre- and post-expansion, with representative plots and aggregate data (n=12). Paired T test results (*p < 0.05) are shown. **F)** Flow cytometric analysis quantifies aNK and cNK cells across TIL clones from 6 patients and 20 clones. Pie charts depict NK cell phenotypes pre- and post-expansion (n=4). In this figure, accumulative data are shown as mean ± SEM.

We further characterized the NK cell receptor expression profile. aNK cells exhibited an activated profile dominated by the expression of CD16, NKG2D, NKp44, and TIGIT, compared to cNK cells that expressed predominantly checkpoint proteins TIM3 and TIGIT. After one month expansion, aNK cells expressed activating receptors NKG2D, NKp30, NKp44, DNAM1, and NKp46. Contrary, cNK cells mostly expressed inhibitory NKG2A (**Figure 4F**).

### aNK cells exhibit greater cytotoxicity against autologous ovarian tumors

Next, we assessed the tumor reactivity of aNK and cNK cells through various functional assays, including effector cell cytotoxicity, pro-inflammatory cytokine production (TNFα and IFNγ), degranulation status (CD107a), and proliferation (Ki67), upon restimulation with autologous tumor cells. We observed that the cultures with the highest tumor cell killing frequency positively correlated with increased amounts of aNK cells. This was not observed with the counterparts cNK and T cell cultures (**Figure 5A-5F**). Additionally, we found a positive correlation between autologous tumor killing and aNK cell IFNγ and TNFα production, moderate correlation with degranulation, but no significant correlation with proliferation (**Supplementary Figure 4B**). Notably, the production of TNFα by aNK cells upon restimulation with autologous tumor cells showed robust correlations with IFNγ production, degranulation, and tumor cell elimination. In contrast, only a weak association was detected between proliferation and tumor cell elimination. These findings strongly reinforce the idea that aNK cells exhibit complete functional proficiency within the TME and underscore their capability for targeted anti-tumor activity.

**Figure 5.**
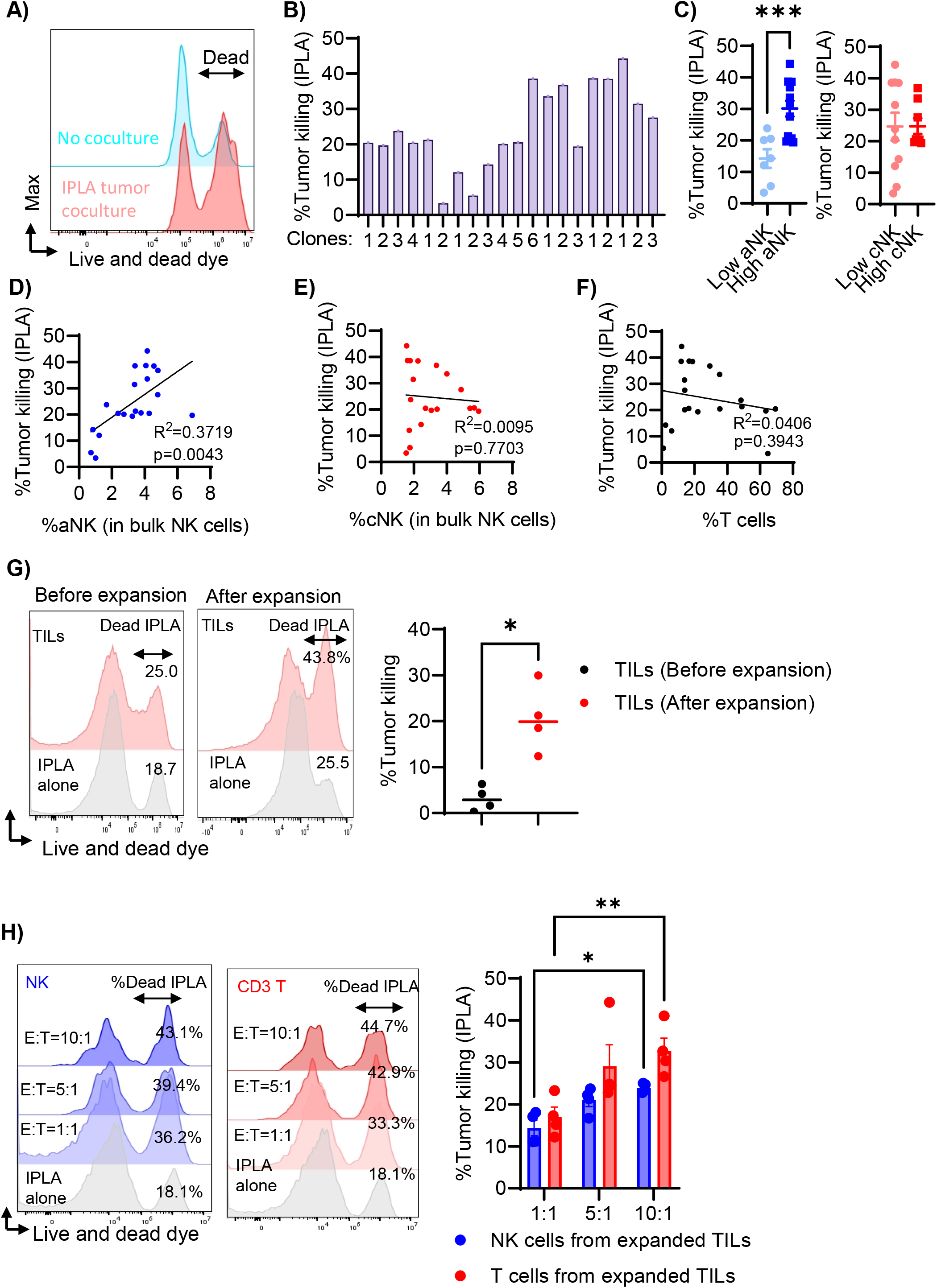
Tumor infiltration lymphocytes (TILs) from ovarian tumors are expanded including aNK cells and cNK cells. **A)** The histogram shows tumor cell death by TIL activity compared to baseline cell death in IPLA tumor cells alone. Tumor killing was measured after a 6-hour interaction between IPLA tumor cells and various TIL clones (n=20). **B)** Analysis of tumor cell death is categorized by TIL clones with high (>3%) or low (<3%) presence of aNK cells (13 and 7 clones, respectively, on the left) and cNK cells (8 and 11 clones, respectively, with >3% or <3% frequency, on the right). Statistical significance was determined using unpaired T tests, with ***p < 0.001 indicating significant differences. **D-F)** Linear regression plots depict the relationship between the frequency of aNK cells (n=20), cNK cells (n=19), and T cells (n=20) versus tumor-killing efficacy. **G)** The impact of TIL expansion on IPLA tumor cell killing was assessed before and after expansion, with significance denoted by *p < 0.05. H) Representative histograms of IPLA tumor killing by isolated NK cells and T cells at different ratios of effector cells: tumor cells (E:T) (Left). Efficiency of TILs in eradicating IPLA tumor cells (n=4 for each group), with significance denoted as *p < 0.05, **p < 0.01. In this figure, accumulative data are shown as mean ± SEM.

Given the observed correlation between intratumoral aNK cell presence, functionality, and tumor killing, we sought to conduct specific tumor killing assays to validate NK cell specific killing. Nevertheless, low numbers of cytotoxic immune cells pre-expansion did not allow for isolating enough NK and T cells. First, comparing pre- and post-expansion, autologous tumor cell killing increased after TIL expansion, suggesting regained cytotoxic capacity through *ex vivo* activation (**Figure 5G**). We next performed the direct tumor killing assay using bead-isolated NK or T cells after expansion. Following expansion, both NK and T cells exhibited the ability to kill autologous IPLA tumor cells, confirming NK cell specific recognition and killing of autologous tumors (**Figure 5H**). Subsequently, we explored whether a specific aNK cell receptor profile correlated with increased tumor killing. We found that aNK cell DNAM-1 expression positively correlated with autologous tumor killing, whereas a strong inverse correlation was observed with the expression of checkpoint receptors, including NKG2A, TIM3, and TIGIT (**Supplementary Figure 4C-4E**). Our data suggest that aNK cells maintain an immunologically active state with a low checkpoint protein profile even within desmoplastic tumors, enabling them to resist TME immunosuppression.

### HD aNK cells can form immunological memory to IPLA tumors

Next, we investigated the ability of healthy blood donor (HD) aNK cells to develop memory responses against IPLA primary tumors. Employing our established DC priming protocol, NK cells were cultured for a period of three to four weeks. Remarkably after three weeks of coculture, aNK cells exhibited the capacity to generate antigen-specific cytokine production, degranulation and proliferation reactions to IPLA tumor cells. Contrasting, cNK cells displayed a stochastic response toward both IPLA and CFPAC tumor cells, characterized by variable cytokine production, degranulation and proliferation (**Figure 6A Supplementary Figure 5A**). However, cNK cell responses became inconsistent by the fourth week of culture. In stark contrast, aNK cells exhibited a consistent and specific reaction towards IPLA cells, maintaining their s p eci f i c target response throughout the entire four-week coculture period with IPLA-lysate loaded DCs (**Supplementary Figure 5B, 5C**).

**Figure 6.**
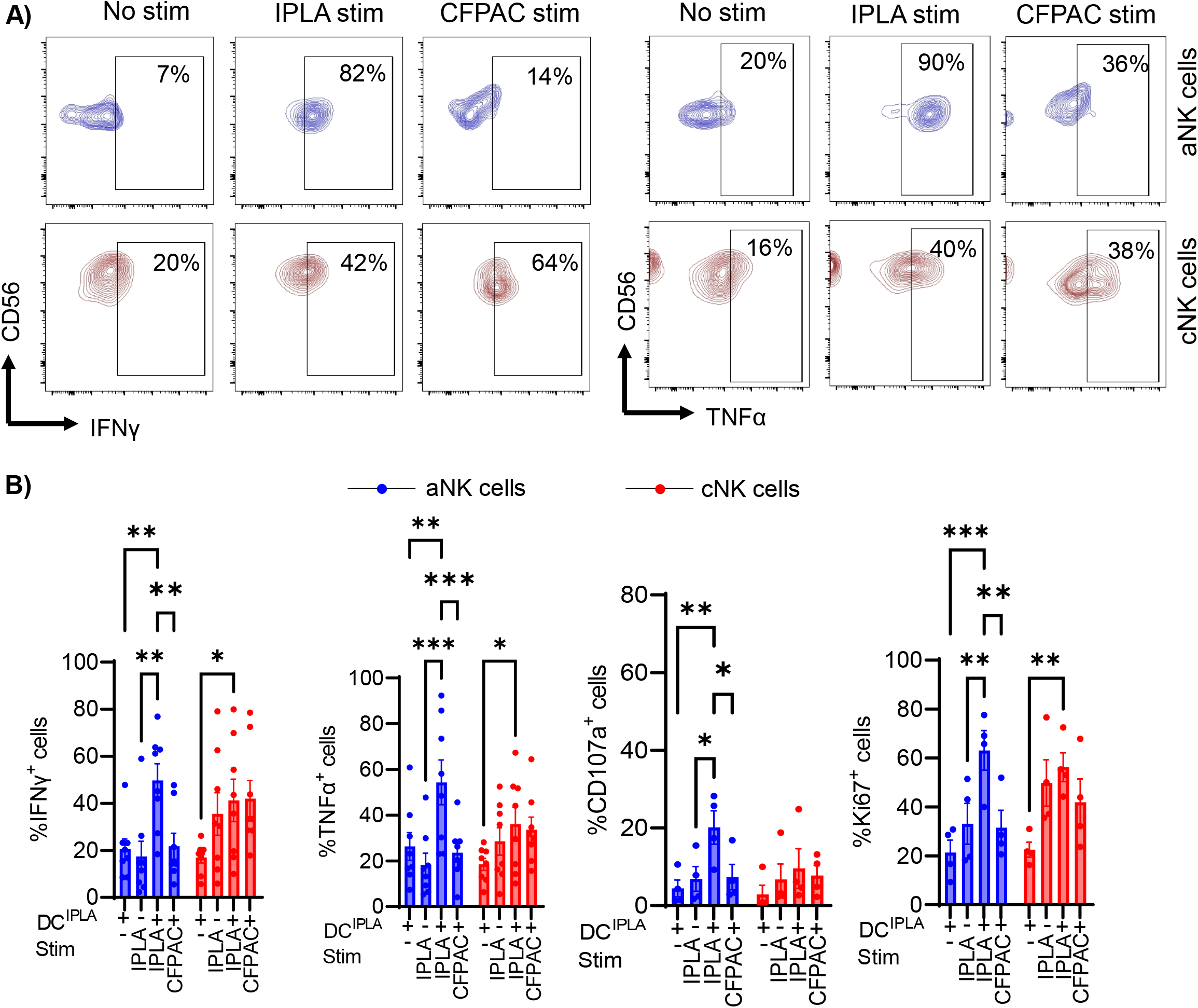
aNK cells have specific recall responses targeting the primary ovarian tumor cells. **A)** Following a three-week co-culture period of NK cells with DCs, re-exposure to IPLA primary tumor cells was conducted. Displayed are representative contour diagrams showing IFNγ^+^ and TNFα^+^ cells within both aNK and cNK cells under various conditions (sequentially from left to right: unstimulated post-DC co-culture, re-stimulated with autologous IPLA cells, and re-stimulated with allogenic CFPAC cells). **B)** Compiled data illustrate the frequencies of IFNγ^+^, TNFα^+^, CD107a^+^and Ki67^+^ cells across various experimental conditions, with the sample size noted as n=8 for IFNγ and TNFα, and n=4 for CD107a and Ki67. Two-way ANOVA was conducted. Significance levels are indicated as follows: *p < 0.05, **p < 0.01, ***p < 0.001. In this figure, all the accumulative data are shown as mean ± SEM.

### aNK cell memory and migration mechanisms toward autologous tumor cells involve HLA-E/NKG2C axis and CXCR2

To gain better insights into the mechanisms involving the DC-aNK cell axis in anti-tumor immunity, particularly in the context of HGSOC, we tested to block molecules that could potentially influence the interaction between aNK cells and DCs. Based on our analysis using single-cell RNA sequencing results, we targeted the following molecules for blockade: NKG2C, NKG2A, HLA-E, HLA-ABC (MHC-I), and MHC-II, at either the priming phase (day 0, day 7, and day 14) or the secondary stimulation phase (recall phase, day 21/28). Blocking these molecules during the priming phase revealed that anti-NKG2A, anti-MHC I, and MHCII were in part inhibiting both cNK and aNK cell function in form of degranulation and cytokine production. Conversely, blocking NKG2C and HLA-E during the secondary stimulation phase resulted exclusively in abolished aNK cell recall responses (**Figure 7A, 7B, Supplementary Figure 6**). These findings suggest that NKG2A, MHC I, and MHCII play crucial roles in educating and activating NK cells to non-self-antigens, whereas HLA-E and NKG2C are pivotal for antigen recognition and subsequent recall responses.

**Figure 7.**
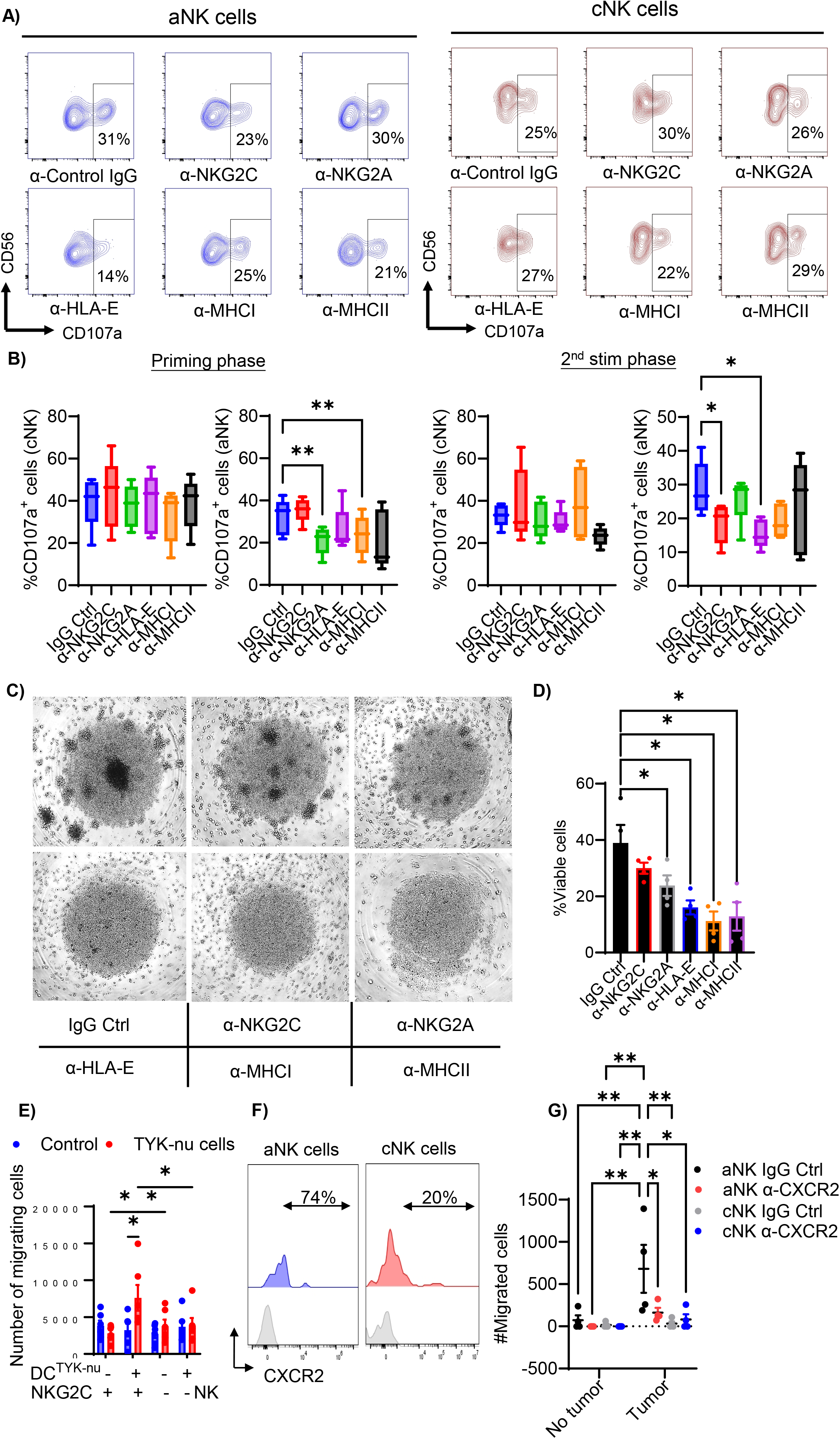
NKG2C and HLA-E are integral to the recall response mechanisms in aNK cells, whereas CXCR2 facilitates the migration of aNK cells to autologous tumors. **A)** Contour diagrams show CD107a^+^ prevalence in aNK and cNK cells after second-phase stimulation blocking with various neutralizing antibodies upon re-exposure to IPLA tumor cells. **B)** Statistical summaries indicate CD107^+^ cNK and aNK cells after priming phase blocking on the left and second-phase restimulation blocking on the right during re-exposure to IPLA tumor cells (n=5), analyzed by one-way ANOVA (*p < 0.05, **p < 0.01). **C)** Representative images display NK clusters and proliferation status under different priming phase blockings with various neutralizing antibodies at week 3. **D)** Statistical summary shows NK cell viability under different priming phase blockings at week 3 (n=5), with data presented as mean ± SEM. **E)** Transwell assay data demonstrate the migration of bead-isolated aNK and cNK cells toward autologous tumor cells over 6 hours under varying conditions (n=6), analyzed using two-way ANOVA (*p < 0.05). **F)** A representative histogram of CXCR2 expression before expansion in aNK and cNK cells from TILs. **G)** Statistical analysis illustrates the migration of aNK and cNK cells from TILs toward autologous IPLA tumors upon treatment with a CXCR2 neutralizing antibody across five samples (n=5), analyzed via one-way ANOVA (*p < 0.05, **p < 0.01, ***p < 0.001). In this figure, accumulative data are shown as mean ± SEM.

Subsequently, we examined NK cell proliferation over three weeks of co-culture in the presence or absence of blocking antibodies. All blocking antibodies reduced NK cell viability, with the exception of anti-NKG2C, which appears to be particularly important for recall responses. NK cells either formed fewer clusters or failed to form clusters altogether, with viability following a similar trend (**Figure 7C, 7D**). This suggests that aNK cells obtain survival/proliferating signals from DCs through these receptors.

Given that in our HGSOC tissues spatial analyses, we observed that aNK cells exhibited a greater capacity to infiltrate tumor areas (**Figure 3A, 3B**), and in our single-cell transcriptomic analyses indicated the potential involvement of CXCR2 in aNK cell-DC interaction intratumorally (**Figure 2F**), we evaluated the migration capacity toward previously primed antigen (TYK-nu) and subsequently blocked CXCR2 in TILs migrating toward autologous tumors. A higher frequency of antigen-primed aNK cells (referred to as NKG2C+) migrated toward TYK-nu cells compared to unprimed aNK cells and both primed and unprimed cNK cells (**Figure 7E**). Additionally, we observed a higher frequency of aNK cells expressing CXCR2 in TILs compared to cNK cells. Upon blocking CXCR2 in migration assays toward autologous IPLA tumors, aNK cell migration was significantly diminished (**Figure 7F**), underscoring the essential role of CXCR2 in aNK cell migration toward tumors.

## Discussion

In this study, we have shown that aNK cells are infiltrating the ovarian TME and are located in the tumor nest. Transcriptome analyses from ovarian cancer patients demonstrated functional aNK cells that interact with DCs. In addition, the function of *ex vivo* expanded aNK cells from ovarian tumor biopsies were significantly associated with high activity and specific killing of autologous tumor cells, suggesting an important function of the intratumoral aNK cells during ovarian carcinoma progression. To date, the function of aNK cells in solid tumors have remained poorly described, although the favorable clinical properties of aNK cells in human hematological malignancies and their contribution to relapse protection have been suggested (*15, 26–30*). For instance, a leukemic murine xenograft model with adoptively transferred cytokine-preactivated memory-like NK cells presented a reduced leukemia burden and an improved overall survival compared to adoptive cNK cell transfer (*31*). Moreover, the high memory-like NK cell cytotoxicity, cytokine production, and persistence *in vivo* were associated with melanoma growth inhibition in mice (*32*). Together with the current data, these studies imply that aNK cells display instrumental anti-tumor properties compared to their counterpart cNK cells in both solid and hematological malignancies, which could be further explained by multiple complementary mechanisms showed in this study.

We found that aNK cells compared to cNK cells in HGSOC display unique phenotypical hallmarks that may enable them to resist the immune escape mechanisms exploited by cancer cells, including down-regulation of the inhibitory receptor NKG2A, and up-regulation of the activation receptors NKG2D, NKp30, NKp44, NKp46, DNAM1, and CD16. The NKG2A receptor confers an inhibitory checkpoint for NK cells (*33*). The anti-NKG2A checkpoint inhibitor has been now tested in a phase II clinical trial supporting the anti-tumor immunity in both by NK cells and T cells and resulting in progression free survival in patients with metastatic cancers at least for four months (*34*), highlighting the importance of the NKG2A immune evasion axis. Malignant ovarian cells express HLA-E, and upon tumor-NK cell interaction, they inhibit NK cell cytotoxic response through HLA-E engagement with the NK-inhibitory receptor NKG2A (*35–37*). Markedly, a very low fraction of aNK cells express NKG2A, potentially enabling their escape from such tumor-immunoevasive mechanisms.

A hallmark of memory T cells is their ability to expand during a secondary encounter with the same antigen, and having a distinct TCR signaling, effector and proliferative potential (*38, 39*). Here, we found that, similar to memory T cells, aNK cell exhibit an intrinsic immunological memory capacity by recognizing tumor antigens following DC priming and perform specific tumor killing and a clonal-like expansion. Notably, TIL clones with higher aNK cell frequencies mediated autologous tumor cell-killing through TNFα and IFNγ secretion, which was not detected in cNK cells. This activity was also observed towards the cell line TYK-nu, which could have tumor-associated antigen similarities to the human primary IPLA cancer cells, considering this cell line originates from a human HGSOC (*40–42*). To date, several studies have reported how aNK cells can recognize specific viral-peptides, loaded on target cell MHC-I, HLA-E and IgG molecules, and undergo clonal-like expansion towards CMV or other viruses, resembling to anti-viral adaptive responses (*43–46*). On the other hand, there is a big knowledge gap about aNK cells in desmoplastic tumors. In our study, we have successfully demonstrated the indispensable role of MHC-I, NKG2A, and MHC-II in the process of tumor-antigen priming for aNK cells, which plays a crucial role in NK cell education as well. Our hypothesis suggests that these molecules play a pivotal role in discerning between self and non-self-peptides. On the other hand, we found that HLA-E and NKG2C are important for the recall responses in aNK cells, which has been earlier reported by us and others in defense against viruses (*12, 47*). Moreover, we found e.g., a greater TNFα secretion by aNK cells following a secondary stimulation with specific tumor cells, which facilitates the activation of the transcription factor AP-1 that is activated both in antigen specific T cells and clonal expanded NK cells (*46, 48*). We also observed a downregulation of CD62L expression in aNK cells following encounter with tumor cells resembling effector memory T cells that downregulate CD62L upon leaving the lymphoid tissues, when homing to peripheral tissues and the site of infection (*38*). Altogether, our data suggest that intratumoral aNK cells exert memory T cell-like properties.

To gain a better understanding of aNK cell immune memory acquisition towards tumor-antigens, we set up an *in vitro* model mimicking DC-mediated T cell activation. Our results indicate that aNK cells possessed recall responses to viable tumor cells following DC tumor-antigen presentation. Whereas previous studies have highlighted DC-NK cell bi-directional crosstalk during immune responses (*49–52*), aNK cell-DC link with immune memory acquisition has not been reported before. To achieve an efficient immune response against tumors, as well as viral-infected cells, DCs are known to promote NK cell function through cytokine secretion. For instance, IL-12 induces NK cell IFNγ-mediated secretion (*53*), IL-15 promotes NK cell development and survival (*54*), and NK cytotoxicity is boosted by type I IFN (*55*). Moreover, intratumoral NK cells can form stable conjugates with DCs, which encourage DC migration and intratumoral abundance through the production of the chemoattractants CCL5, XCL1, and XCL2 (*56, 57*), thus hampering tumor growth and development altogether. Markedly, the scRNA-seq analysis highlighted the chemoattractant XCL1 to be differentially expressed by intratumoral aNK cells. These studies have demonstrated that NK cells and DC encounters are physiologically relevant and in line with our findings that aNK cells and DC interaction fostered aNK cell immunological memory.

In our rapid expansion protocol, NK cells were able to significantly expand, however it should be taken into account that there was no selected expansion for aNK cells. Although ongoing new insights into epigenetic and transcriptomic mapping could lead to the targeted use of specific growth factors, to date, the best aNK cell expansion protocols still rely on an initial “seeding population” with large numbers of aNK cells (*58*). A recent expansion protocol has been established utilizing HLA-E with an HLA-G-leader-derived peptide expressing feeder cells, to reach >90% pure aNK cells that express single-killer-cell immunoglobin-like receptor (KIR)+. Unfortunately, these cells are not antigen specific and rely on HLA-mismatching to effectively kill acute myeloid leukemia cells (*59*). Therefore, to circumvent the outgrowth of undesired immune cell subsets, more optimal/reliable protocols should be developed taking into account antigen presentation, multiple cytokine combinations, and sequential dosing schedules based on monitorization of the cell population frequencies and function. In our model, expanded cells were exposed to saturating cytokine and serum concentrations for long periods of time, which were outside normal physiological ranges. This could greatly promote immune cell proliferation yet terminally differentiate the cells into exhausted/anergic effector phenotypes, or favor expansion of less mature NKG2A^+^CD56^dim^ NK cells expressing polyclonal KIR repertories, as previously reported (*60*). The protocol used in this study, although not specifically expanding aNK cells, does not alter their functionality. Since the ultimate goal of conducting such protocol is to potentially use adoptive aNK cell transfer as an immunotherapeutic approach, future research into achieving high aNK cell numbers with tumor-antigen specificity, optimal *in vivo* functionality and persistence, will aid in the optimization of these protocols.

The data presented here suggest that the unique properties of aNK cells could be harnessed in cancer NK cell-based therapy. The results also show the potential of DCs to enhance aNK cell function and tumor-cell reactivity, which could lead to interesting alternative fields of research for improving antitumor immunotherapy, such as vaccine development or induced pluripotent stem cell (iPSC)-derived NK cell generation (*61*). DC-based cancer vaccination targeting tumor-antigens have proved to be biologically safe and effective (*62*). In conclusion, the significance of these findings is in providing novel insights into aNK cell antigen specificity and recall responses for the first time, adding the DC-NK cell axis to potential therapeutic strategies against aggressive and metastatic desmoplastic tumors.

## Acknowledgements

We would like to thank and Wenche S. Prestvik at the Department of Biomedical Laboratory Science, Norwegian University of Science and Technology (NTNU), for helping to coordinate the tissue sectioning. Cutting the FFPE tissue sections was performed at the Cellular and Molecular Imaging Core Facility (CMIC), NTNU and Histology Core Facility, Karolinska Institute. And our study is funded by the following funding: National Research Foundation, NRF-CRP26-2021RS-0001 (LKP and SYN), Swedish Cancer Society 211888Pj (KL); Norwegian Cancer Society 216113 (KL); Karolinska Institutet doctoral grant 2–5586/2017 (KL/OG), KI Stiftelser och Fonder, 2020-01829 (DS); Swedish Cancer Society 200169F (DS); Swedish Cancer Society 201128Pj (DS); China Scholarship Council 201906280459 (YS); Stiftelsen Clas Groschinskys Minnesfond M2258 (DS).

## Competing interests

Authors declare that they have no competing interests.

## Notes

### Competing Interest Statement

The authors have declared no competing interest.

