## Supplementary material for "CXCR2-mediated recruitment of adaptive NK cells with NKG2C/HLA-E dependent antigen-specific memory enhances tumor killing in ovarian cancer"

**Supplementary Figure 1 related to Figure 1**

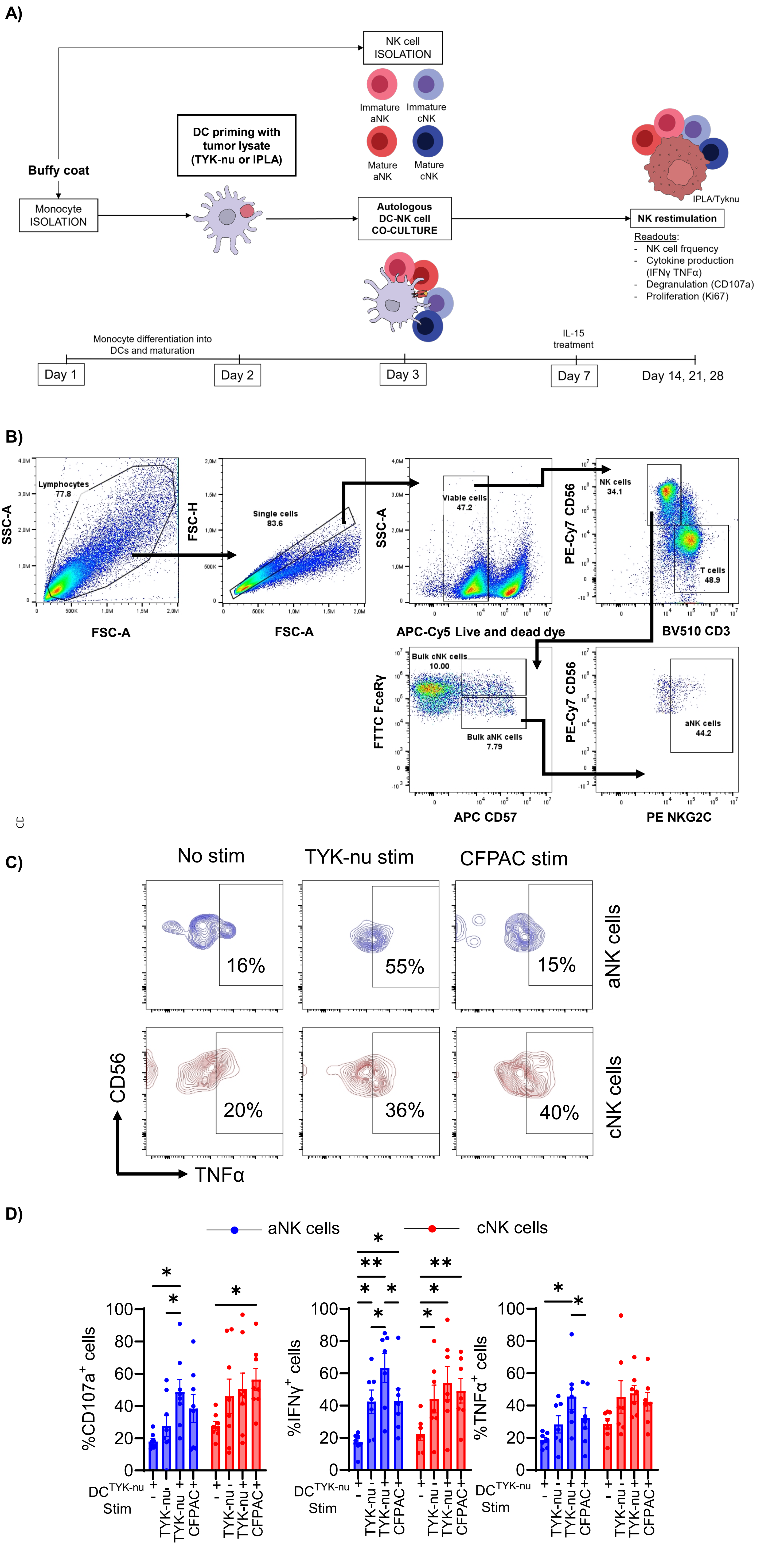

**Supplementary Figure 2 related to Figure 1**

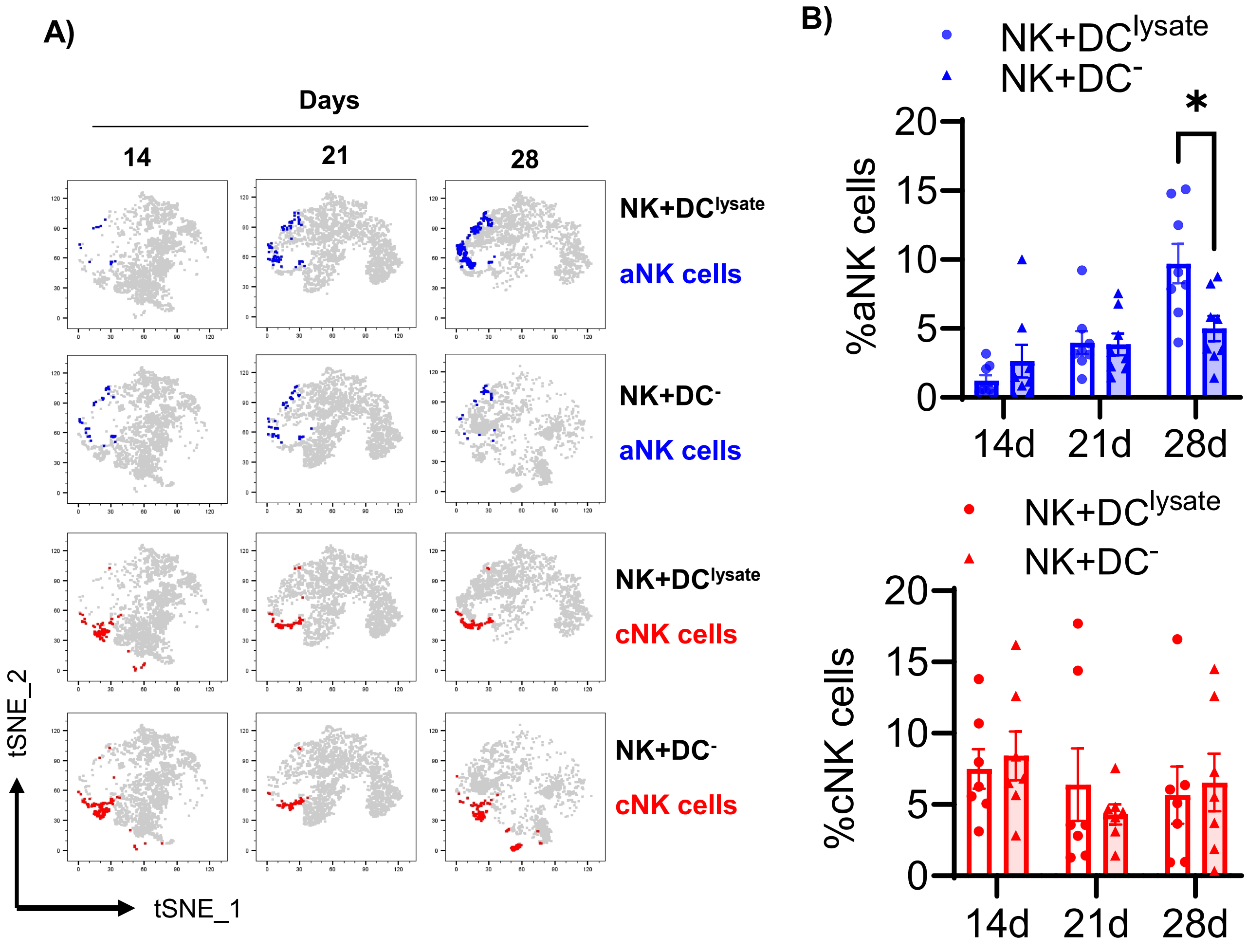

### Supplementary Figure 3 related to Figure 2

A)

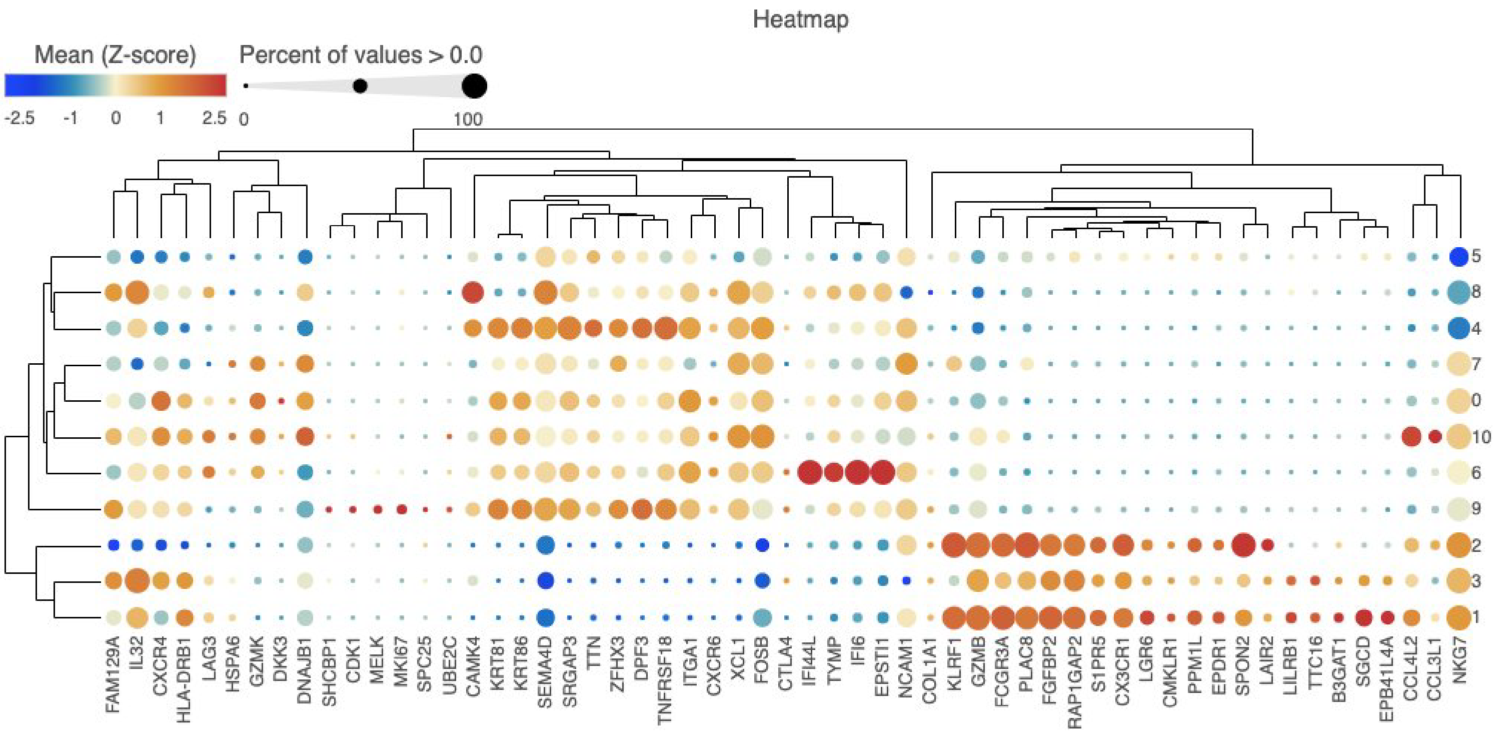


B)

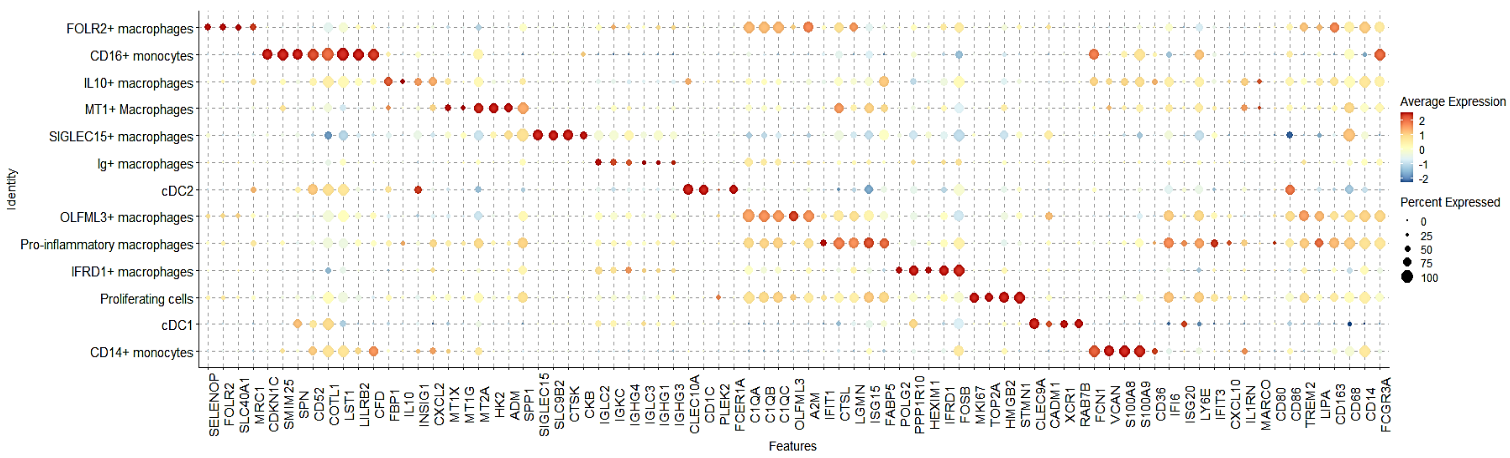

**Supplementary Figure 4 related to Figure 4 and Figure 5**

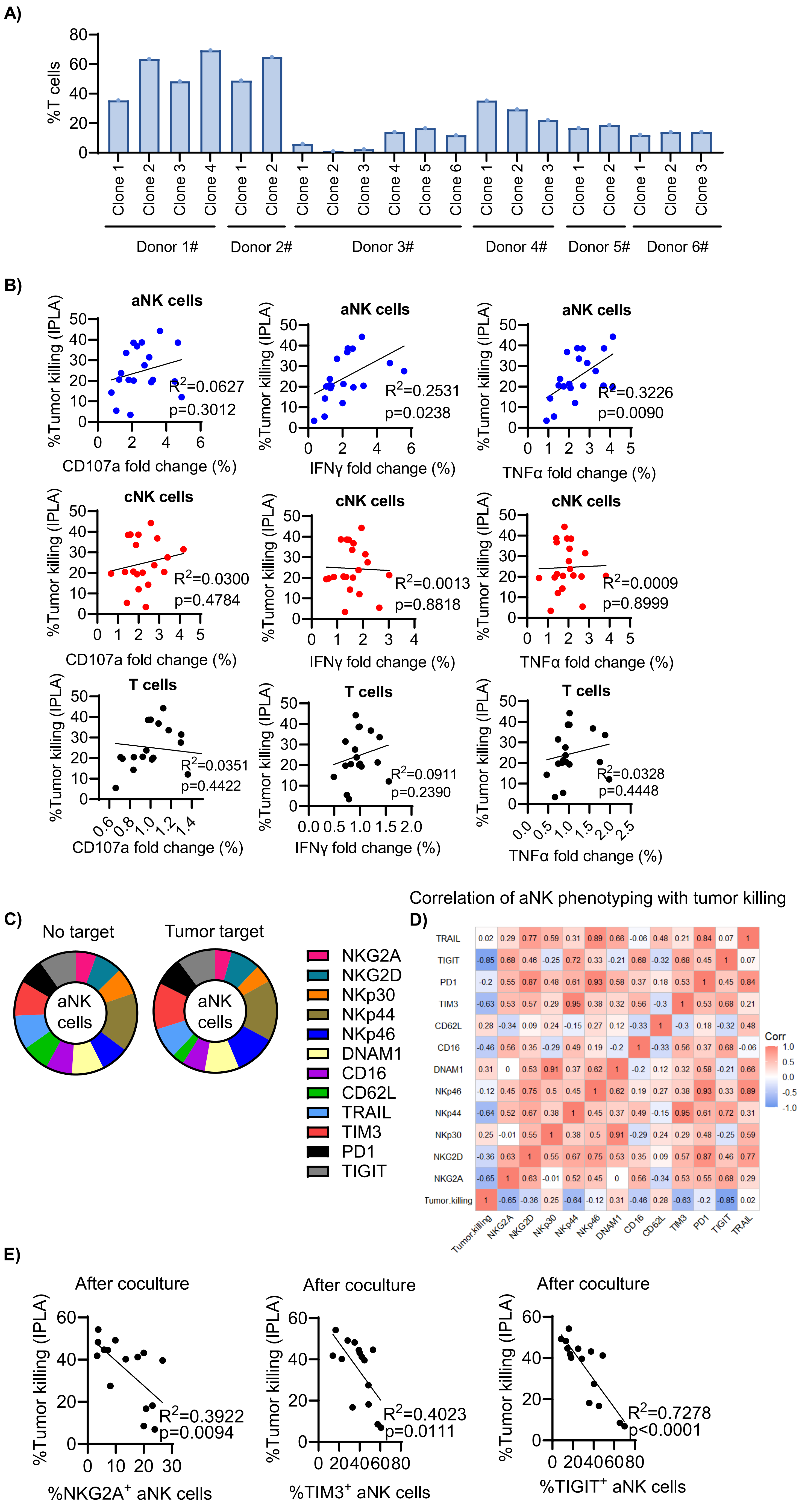

**Supplementary Figure 5 related to Figure 6**

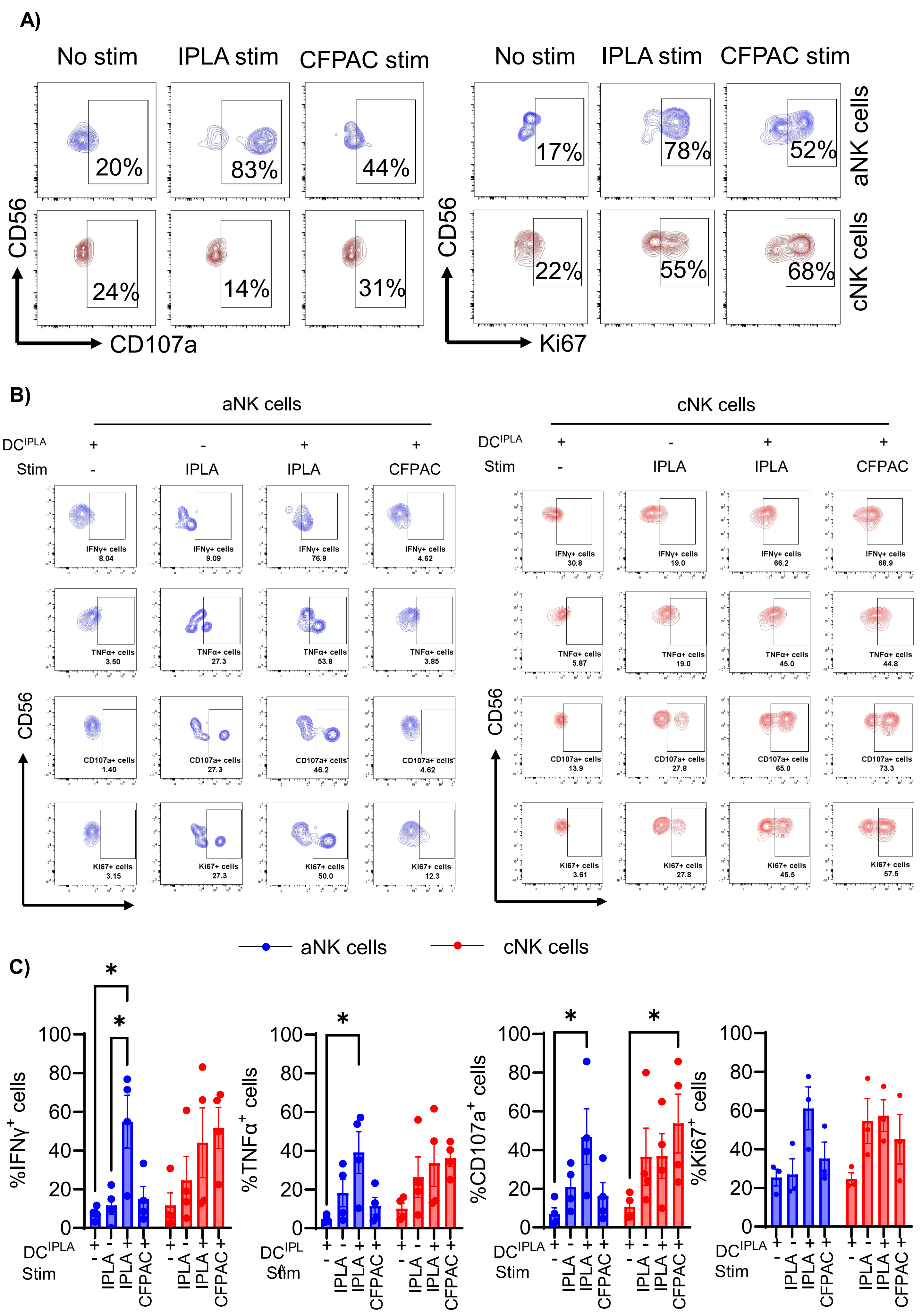

**Supplementary Figure 6 related to Figure 7**

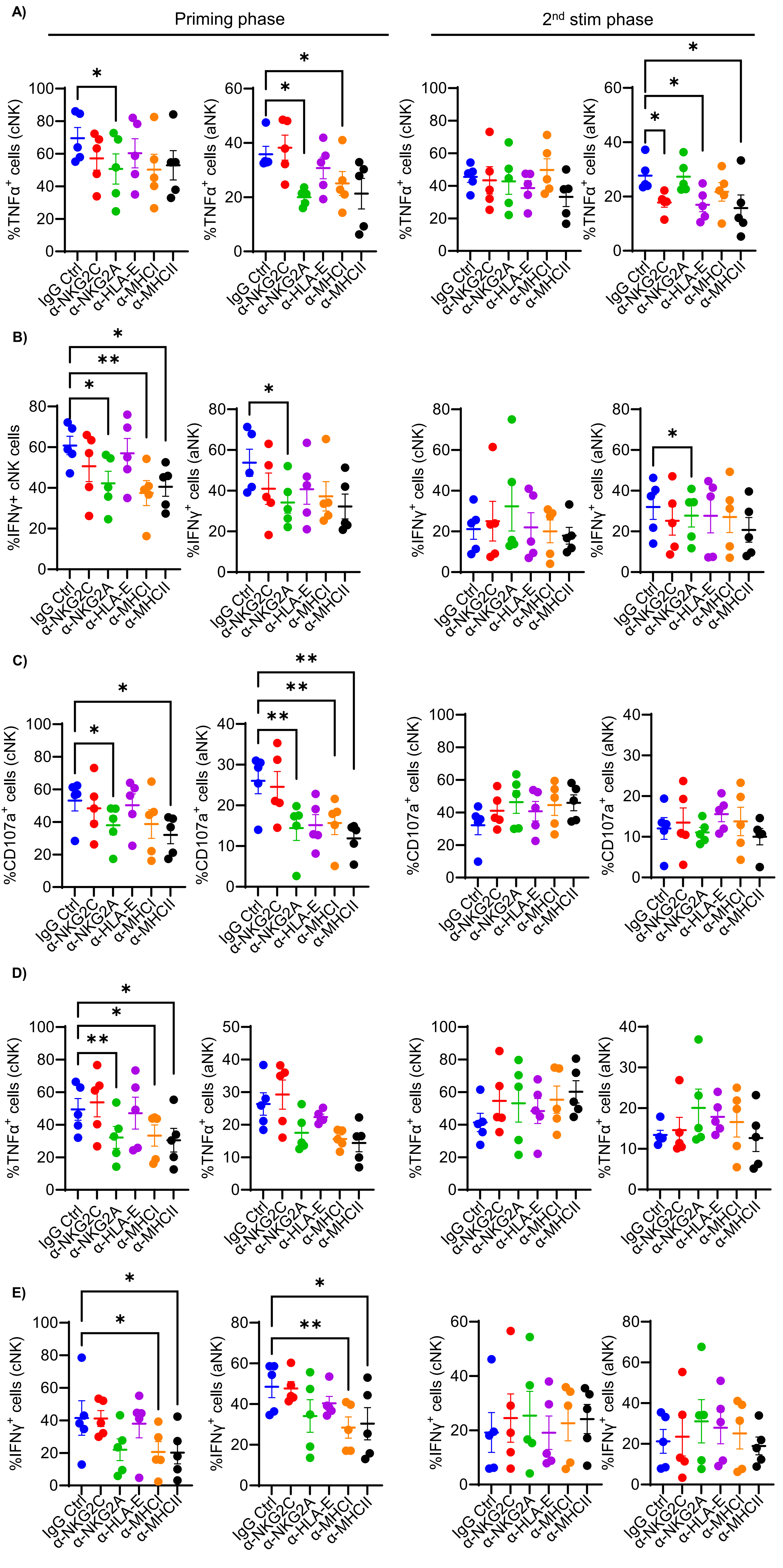

**Supplementary Fig 1. Workflow for the NK cells restimulation and gating for aNK and cNK cells**

**A)** NK cells and monocytes were isolated from buffy coats. Monocytes were then used to cultivate dendritic cells (DCs), which were subsequently primed with tumor lysates or left unprimed. These DCs, either loaded with tumor lysates (TYK-nu or IPLA primary tumor lysates) or left unloaded, were co-cultured with NK cells. The NK cells underwent restimulation with either autologous or allogeneic tumor cells at intervals of the 3rd, and 4th week. **B)** The gating strategy was employed to distinguish between aNK and cNK cells: aNK cells were identified as $CD3^{-}CD56^{+}CD57^{+}Fc\varepsilon R\gamma^{-}NKG2C^{+}$, whereas cNK cells were identified through the $CD3^{-}CD56^{+}CD57^{+}$ $Fc\varepsilon R\gamma^{+}$ markers. **C)** At 3rd week, the representative contour plot of $TNF\alpha^{+}$ cells among aNK and cNK cells is shown under different states (From left to right: no stimulation after DC coculture, TYK-nu restimulation as autologous restimulation, CFPAC restimulation as allogenic restimulation). **D)** After 4 weeks coculture, frequencies of $CD107a^{+}$ cells (n=7), $IFN\gamma^{+}$ cells (n=7), $TNF\alpha^{+}$ cells (n=7) under different conditions: $DC^{TYK\text{-}nu}$ +TYK-nu as autologous restimulation; $DC^{TYK\text{-}nu}$+CFPAC as allogenic restimulation; $DC^{-}$ (without TYK-nu lysates) + TYK-nu as first stimulation, both aNK cells (blue) and cNK cells (red) are shown. Two-way ANOVA test was performed for all cumulative data. Data are presented as mean ± SEM of the mean (SEM), and p values are represented as $*p < 0.05$, $**p < 0.01$, $***p < 0.001$.

**Supplementary Fig 2. aNK cells from bulk NK cells can expand upon DC coculture**

**(A-B)** We illustrate both the individual plots and the aggregated quantitative analysis of aNK cells, alongside cNK cells in coculture with DCs, represented in blue and in red respectively (n=8). Representative tSNE maps and cumulative data of frequencies of aNK and cNK over an interval ranging from two to four weeks are shown. Statistical analysis was performed, data is shown as mean ± SEM, and p-values are represented as $*p < 0.05$.

**Supplementary Fig 3. Marker genes from differentially expressed genes (DEGs) feature the subclusters of NK cells and myeloid cells**

**A)** A heatmap that displays the marker genes from DEGs for the 11 clusters of NK cells is presented. **(B)** A heatmap that displays the marker genes from DEGs for the 13 clusters of myeloid cells is presented.

**Supplementary Fig 4. Intratumoral T cell frequency and functionality fail to correlate with the tumor killing**

**A)** T cells were quantified by flow cytometry for each TIL clone selected from the independent IPLA patients, n=6 patients and n=20 TIL clones. **B)** Linear regression analyses of aNK, cNK, and T cell functionality with IPLA cell killing respectively. **C)** The pie charts display the phenotypes of aNK cells before and after coculture with autologous tumor cells. **D)** The following aNK cell (n=15) receptors were analyzed NKG2A, NKG2D, NKp30, NKp44, NKp46, DNAM1, CD16, CD62L, TIM3, PD1, TIGIT and TRAIL, and correlation heatmap is shown indicating the correlations between the phenotype of aNK cells and tumor-killing. **E)** The linear regression analyses of the frequency of NKG2A$^{+}$, TIM3$^{+,}$ and TIGIT$^{+}$ aNK cells after coculture with the autologous tumor killing (n=15) are respectively shown.

**Supplementary Fig 5. aNK cells preserve the recall responses toward IPLA at 4$^{th}$ week**

**A)** Representative plots of the frequencies of CD107a$^{+}$ cells and Ki67$^{+}$ cells in aNK cells and in cNK cells following restimulation with autologous (IPLA) and allogenic (CFPAC) tumor cells after three weeks coculture with DCs with or without IPLA tumor lysates. **B)** Following a four-week co-culture, we assessed the prevalence of IFNγ$^{+}$ (n=4), TNFα$^{+}$ (n=4), CD107a$^{+}$ (n=4), and Ki67$^{+}$ cells (n=3) under different conditions. **C)** To analyze the aggregate data, we utilized a two-way ANOVA test. The findings are depicted as the mean ± SEM, showcasing all p-values <0.1. Observations with p-values > 0.1 were omitted from the depiction.

**Supplementary Fig 6. NKG2A, MHC-I, and MHC-II are crucial for primary NK cell responses, while HLA-E and NKG2C are essential during recall responses to IPLA tumors**

**A, B)** Cumulative data showing TNFα$^+$ and IFNγ$^+$ cNK and aNK cells when subjected to primary and secondary stimulation with IPLA tumor cells (n=5) in the presence of blocking antibodies stated either during the primary (left) or during the re-exposure (right) phase. Statistical analyses were performed using One-way ANOVA, $*p < 0.05$, $**p < 0.01$. **C-E)** Cumulative data showing degranulation (CD107a), TNFα$^+$, and IFNγ$^+$ in cNK and aNK cells when subjected to primary and secondary stimulation with CFPAC tumor cells (n=5) in the presence of blocking antibodies stated either during the primary (left) or during the re-exposure (right) phase. Statistical analyses were performed using One-way ANOVA, $*p < 0.05$, $**p < 0.01$.
